# Targeting a central feature of asthma using a cell type-selective IL-13-responsive enhancer

**DOI:** 10.1101/2021.10.08.463596

**Authors:** Kyung Duk Koh, Luke R. Bonser, Walter L. Eckalbar, Jiangshan Shen, Ofer Yizhar-Barnea, Xiaoning Zeng, Dingyuan I. Sun, Lorna T. Zlock, Walter E. Finkbeiner, Nadav Ahituv, David J. Erle

**Author notes:** Equal contributions (Kyung Duk Koh, Luke R. Bonser, Walter L. Eckalbar).

## Abstract

IL-13 is a central mediator of asthma^1–3^. Here, we used genome-wide approaches to characterize genes and regulatory elements modulated by IL-13 and other asthma-associated cytokines in airway epithelial cells and showed how they can be used for therapeutic purposes. Using bulk and single cell RNA-seq, we found distinctive responses to IL-13, IL-17, and interferons in human bronchial epithelial basal, ciliated, and secretory cells. H3K27ac ChIP-seq revealed that IL-13 had widespread effects on regulatory elements. Detailed characterization of an enhancer of *SPDEF*, a transcription factor required for pathologic mucin production, revealed that STAT6 and KLF5 binding sites cooperate to drive IL-13-dependent transcription selectively in secretory cells. Using this enhancer to drive CRISPRi and knockdown either *SPDEF* or the mucin *MUC5AC* showed the potential use of this approach for asthma therapeutics. This work identifies numerous genes and regulatory elements involved in cell type-selective cytokine responses and showcases their use for therapeutic purposes.

---

Cytokine effects on the airways play a central role in the pathogenesis of asthma, a disease that affects ∼262 million individuals worldwide^4^. IL-13-, interferon-, and IL-17-stimulated gene expression is increased in distinct subsets of individuals with asthma, and these cytokine signatures are associated with disease severity, airway inflammatory responses, and responses to therapy^3,5,6^. These phenomena reflect pleiotropic effects of cytokines in the airway. For example, IL-13, which is implicated in at least half of individuals with asthma, induces eosinophilic inflammation^1,2^, increased airway smooth muscle contractility^7^, fibroblast proliferation^8^, and mucus overproduction^9,10^. Using transgenic mouse models, we previously showed that direct effects of IL-13 on the airway epithelium are sufficient to induce two key features of asthma, mucus overproduction and airway hyperreactivity, in the absence of airway inflammation or fibrosis^9^. A more complete understanding of cell type-selective responses to IL-13 and other asthma-associated cytokines could improve our understanding of how these cytokines contribute to asthma and identify new approaches for therapy.

Various mechanisms account for the ability of cytokines to produce different effects on different cells. For example, the type 2 cytokine interleukin-4 (IL-4) utilizes the type 1 IL-4 receptor (IL-4Rα/γc) in lymphocytes and the type 2 receptor (IL-4Rα/IL-13Rα) in many non-hematopoietic cells, and downstream signaling pathways including Jak/STAT, insulin receptor substrate (IRS), and others may also differ between cell types^11,12^. Differences in the state of genomic *cis*-regulatory sequences have also been shown to contribute to cell type-specific responses^13^. Studies in macrophages illustrated that enhancer repertoires established in differentiated cells can bind to transcription factors activated by cytokines, and that cytokine stimulation can also result in activation of latent enhancers in these cells^14^. However, our understanding of how enhancers control cytokine effects on individual genes in other cell types remains limited. Genome-wide methods such as single cell RNA sequencing (scRNA-seq) and chromatin immunoprecipitation followed by sequencing (ChIP-seq) provide the opportunity for detailed investigation of cytokine effects on gene expression and gene regulatory elements. Here, we use these and other methods to characterize transcriptional and epigenetic effects of IL-13, interferons (IFNs), and IL-17 on the airway epithelium, a complex tissue comprised of basal, ciliated, and secretory cells as well as other less common cell types. Our findings identify cell type-specific effects of these cytokines within the epithelium and demonstrate how enhancers that integrate cell type and cytokine signals can be used for synthetic biology approaches that target asthma.

## Results

### Cytokines induce cell type-selective transcriptional responses in HBECs

We used bulk RNA sequencing (RNA-seq) to assess the transcriptional effects of asthma-associated cytokines on primary HBECs from six individuals grown at air-liquid interface (ALI). A portion of the results from this experiment have previously been used to identify IFN-stimulated genes in asthma^6^ and IL-17-stimulated genes in chronic obstructive pulmonary disease^15^. Integrated analysis of the complete dataset revealed 4,965, 4,799, 5,408, and 8,246 genes that were differentially expressed (FDR < 0.1) in response to IFN-α, IFN-γ, IL-17, and IL-13, respectively (Supplementary Data 1). The effects of each cytokine were distinct, and each cytokine had similar effects in all donors (Supplementary Fig. 1). Since effects of IFN-α and IFN-γ were similar (Supplementary Fig. 1e), we did not include IFN-γ in subsequent experiments. Effects of combined stimulation with IFN-α and IL-13 were largely additive rather than synergistic (Supplementary Fig. 1f)

We used scRNA-seq to investigate cell type-specific transcriptional effects of IFN-α, IL-13, and IL-17 in cultures from four donors (Supplementary Fig. 2a-c and Supplementary Data 2). IFN-α produced a similar transcriptional response in each of the major cell types (basal, ciliated, and secretory cells, Fig. 1a, b, and e). In contrast, many effects of IL-13 and IL-17 differed between these three cell types (Fig. 1c, d, and f and Supplementary Fig. 2d-f). To quantify cell type-selective effects, we used a linear model with terms for cell type, cytokine effect, and the interaction between the two. The interaction between IFN-α stimulation and cell type was significant (log_2_ fold-change difference > 1 and FDR < 0.1) for 58 genes in secretory cells compared with basal cells and 52 genes in secretory cells compared with ciliated cells. IL-17 led to cell type-selective effects for 32 and 108 genes in secretory cells compared with basal and ciliated cells, respectively. IL-13 had substantially more cell type-selective effects: 292 and 166 genes in secretory cells compared with basal cells and ciliated cells, respectively. Gene set enrichment analysis revealed that IL-13 affected different cellular processes in these three cell types (Supplementary Fig. 3a and Supplementary Data 3). As expected, IL-13-induced transcripts in secretory cells were highly enriched for a set of goblet cell genes previously defined in an scRNA-seq analysis of epithelial cells from human lung^16^ (Supplementary Fig. 3b).

**Figure 1.**
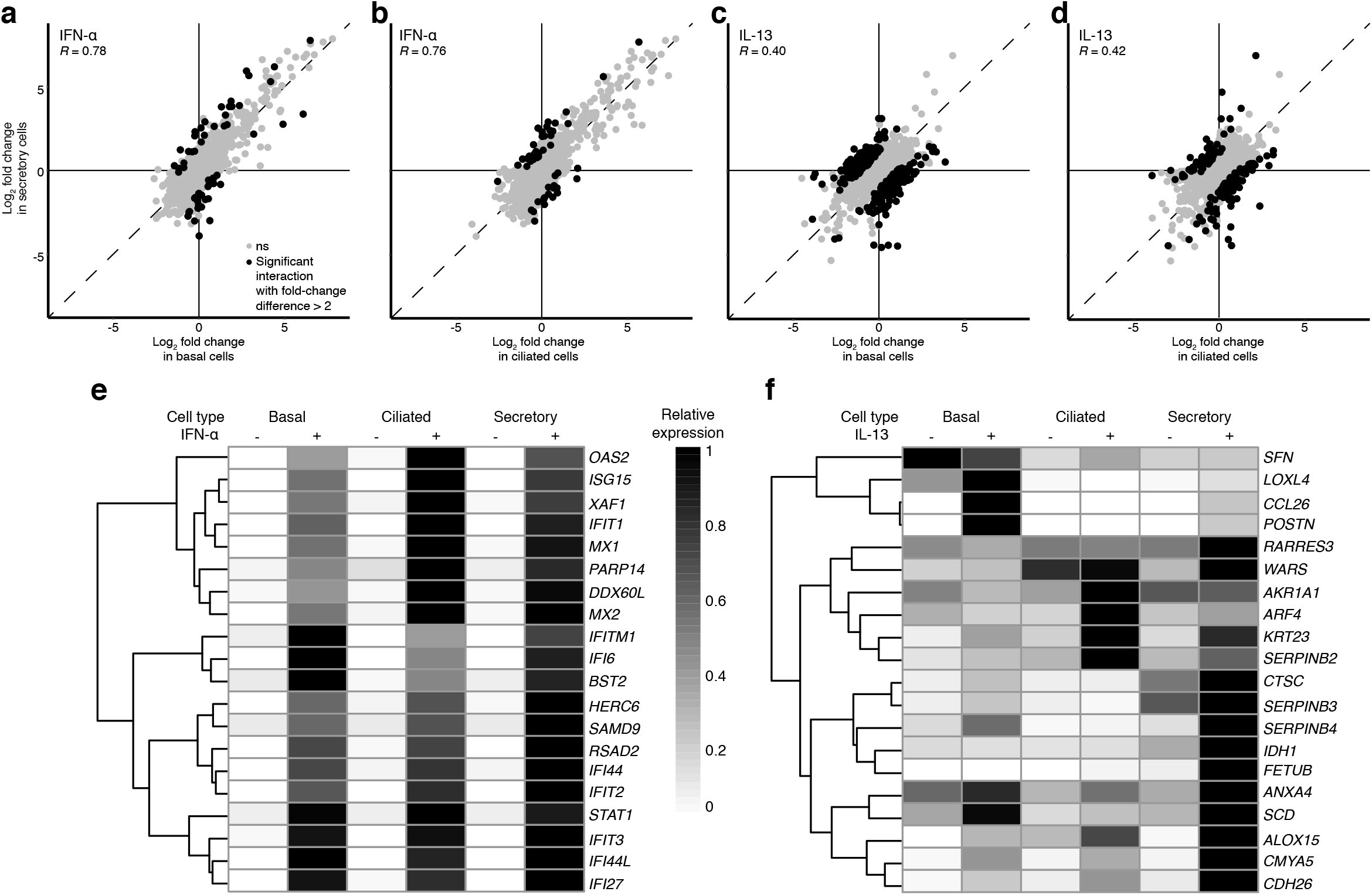
IFN-α and IL-13 induce gene expression changes in each major subpopulation of bronchial epithelial cells. **a**-**d**, Effects of IFN-α (**a** and **b**) and IL-13 (**c** and **d**) on secretory cells compared with basal cells (**a** and **c**) or ciliated cells (**b** and **d**). Each point represents a single gene. Genes that were regulated differently between cell types are in black (log_2_ fold-change difference > 1 and FDR < 0.1 for interactions between cell type and cytokine effect), and all other genes are in grey. *R*, Pearson correlation coefficient (*p* < 2.2 × 10^−16^ for all comparisons). **e** and **f**, Relative expression of the 20 genes that were most strongly regulated by IFN-α (**e**) and IL-13 (**f**). These 20 genes were selected based on the highest absolute fold change in any of the three cell types.

### IL-13 induces epigenomic changes in HBECs

Since enhancers have major roles in determining cell-specific gene expression, we investigated whether cytokine stimulation had effects on enhancer activity. We performed ChIP-seq using an antibody recognizing acetylated lysine 27 on histone 3 (H3K27ac), a marker of active enhancers and promoters^17,18^. Comparison of unstimulated cells with cells stimulated with IFN-α or IL-17 revealed that only 12 of 24,113 (0.05%) or 2 of 29,754 (0.01%) of H3K27ac-associated regions were modulated by IFN-α or IL-17, respectively. In contrast, 602 of 19,629 regions (3.1%) were affected by IL-13 (387 enriched, 215 depleted). Approximately one-third of the IL-13-enriched regions are located in putative promoters (Fig. 2a), suggesting that most IL-13-induced sequences function as distal regulatory elements. Regions enriched in H3K27ac upon IL-13-stimulation were preferentially located near the transcription start sites (TSSs) of IL-13-induced genes; conversely, regions depleted in H3K27ac tended to be located near genes with decreased expression (Fig. 2b). *De novo* motif discovery showed that the most highly enriched motif within IL-13-induced H3K27ac regions conform to a previously described STAT6 motif^14^ (Fig. 2c). Collectively, these data indicate that IL-13-induced transcriptional changes are accompanied by significant changes in H3K27ac levels. In contrast, IFN-α- and IL-17-induced changes in gene expression were largely independent of H3K27ac modifications, suggesting that other regulatory mechanisms drive transcriptional responses to IFN-α and IL-17.

**Figure 2.**
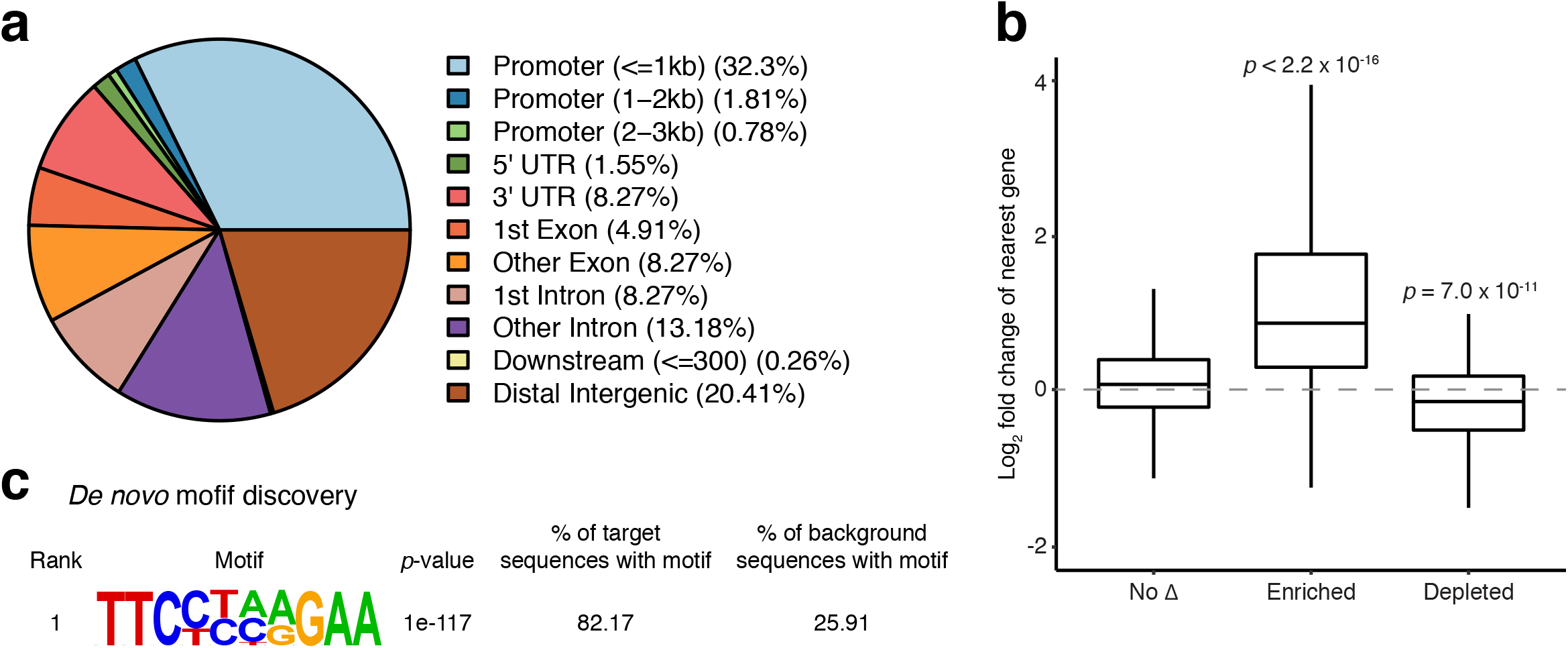
IL-13 induces enrichment of H3K27ac peaks near IL-13-stimulated genes. **a**, Distribution of IL-13-enriched H3K27ac peaks (N = 387) by genomic region classification. **b**, Bulk RNA-seq expression changes in genes nearest H3K27ac ChIP-seq peaks that were not significantly affected (*p* > 0.1) by IL-13 (“No Δ”, 26,875 peaks) or were significantly (FDR < 0.1) enriched (387 peaks) or depleted (215 peaks) following IL-13 stimulation. *p*-values for comparison with unaffected peaks by two-sided Wilcoxon test are shown. **c**, The most highly enriched motif discovered within regions with enriched H3K27ac following IL-13 stimulation closely resembles a previously defined STAT6 binding motif.

### A distal *SPDEF* enhancer plays a role in goblet cell differentiation and mucus hyperplasia

One region with increased H3K27ac following IL-13 stimulation is located 29 kb upstream of the SAM pointed domain-containing Ets transcription factor (*SPDEF*) TSS (Fig. 3a). SPDEF is an IL-13-inducible transcription factor required for airway epithelial cell mucus metaplasia and changes in mucus organization and function^19,20^. Our scRNA-seq results showed that IL-13 stimulation induced *SPDEF* in secretory cells (1.99-fold, FDR = 1.24× 10^−18^). *SPDEF* expression in basal and ciliated cells was >10-fold lower and was minimally affected (<1.1-fold) by IL-13. To determine the function of the putative *SPDEF* enhancer (SPDEFe) in HBECs, we used dCas9-KRAB-based CRISPR interference (CRISPRi, Fig. 3b-e). In cells expressing a control (non-targeting) gRNA, IL-13 induced expression of *SPDEF* and the *SPDEF-*regulated genes *FOXA3* and *MUC5AC*. Targeting *MUC5AC* directly using gRNAs against its TSS reduced *MUC5AC* mRNA but did not affect *SPDEF* or *FOXA3*, which are upstream of *MUC5AC*. In contrast, targeting the *SPDEF* TSS reduced *SPDEF, FOXA3*, and *MUC5AC* mRNA levels as well as MUC5AC protein. Targeting the putative enhancer SPDEFe had very similar effects on *SPDEF, FOXA3*, and *MUC5AC*. Targeting SPDEFe did not affect SPDEF-independent effects of IL-13 and did not affect expression of other genes located near the *SPDEF* locus (Supplementary Fig. 4). For more precise functional mapping of the SPDEFe region, we tested additional gRNAs and found that gRNAs targeting sequences 2-4 kb from SPDEFe had minimal effects on levels of MUC5AC (Fig. 3f and Supplementary Table 1). Taken together, these results indicate that targeting a limited region including SPDEFe effectively and selectively suppresses IL-13-induced *SPDEF* and SPDEF*-*dependent genes.

**Figure 3.**
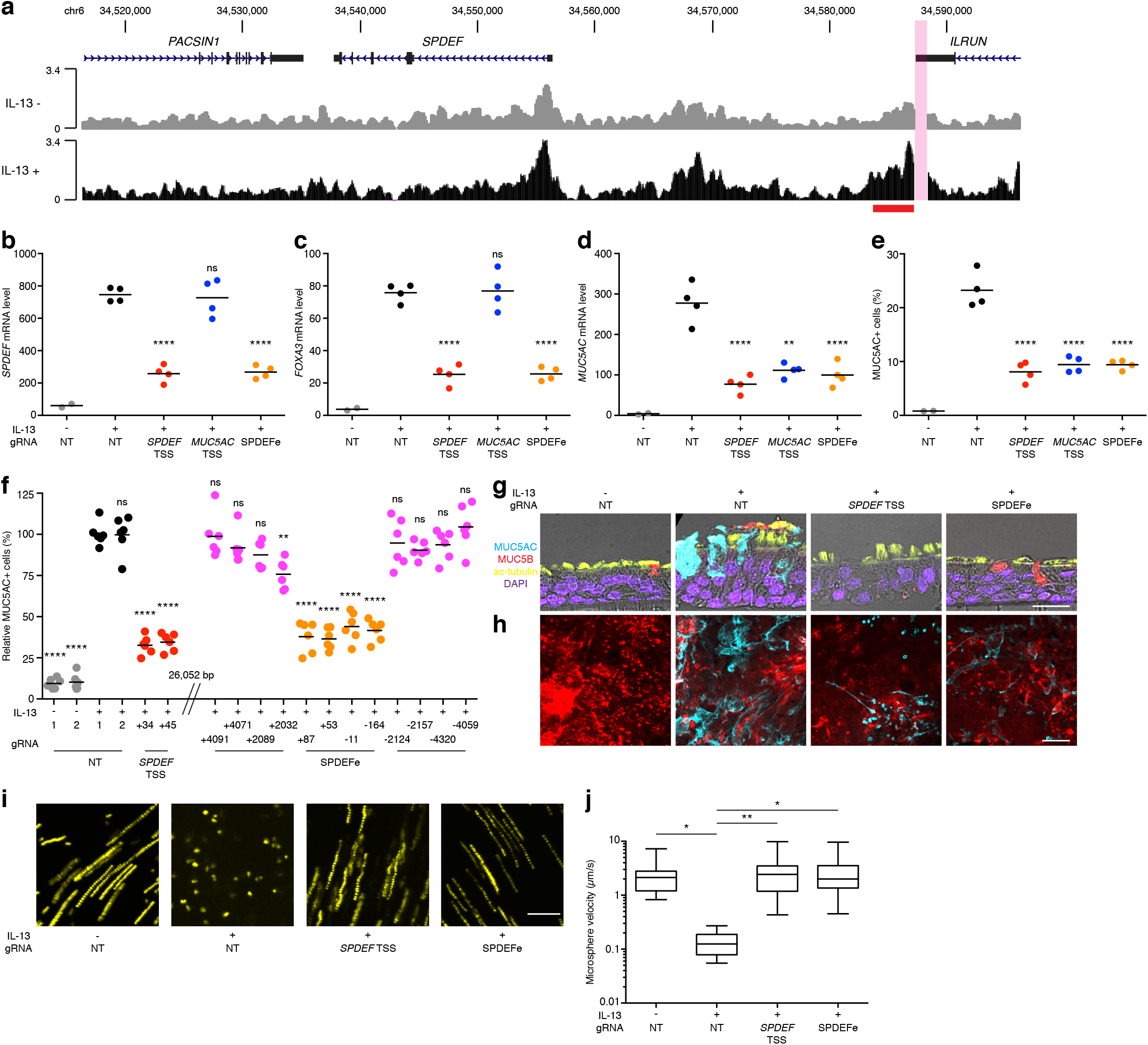
CRISPRi targeting of the *SPDEF* enhancer prevents IL-13-induced mucus metaplasia and mucostasis. **a**, IL-13 induces increased H3K27ac within a region (chr6:34,583,800-34,587,500) 29 kb upstream from the *SPDEF* TSS. Tracks represent H3K27ac ChIP-seq read depth from HBECs cultured with (black) and without (grey) IL-13 stimulation (mean of four donors). The lone peak in this region that was affected by IL-13 (FDR < 0.1) is indicated by a red bar below the tracks. An ENCODE-blacklisted repetitive region that cannot be mapped reliably is highlighted in magenta. **b**-**d**, CRISPRi effects on gene expression. HBECs were transduced with lentiviruses driving expression of *dCas9*-*KRAB* and either non-targeting (NT) control sgRNAs (grey/black) or sgRNAs targeting the *SPDEF* promoter (red), the *MUC5AC* promoter (blue), or SPDEFe (orange). After differentiation, cells were left unstimulated or stimulated with IL-13 for 7 days, as indicated. Expression of *SPDEF* (**b**), *FOXA3* (**c**), and *MUC5AC* (**d**) were measured by quantitative real-time PCR (qRT-PCR). Values are mRNA copy numbers per 1,000 copies of *GAPDH*. Each point corresponds to a different gRNA targeting the indicated region, tested separately in a single culture well from the same donor. ns, not significant; **, *p* < 0.01; and ****, *p* < 0.0001 for comparison with IL-13-stimulated HBECs with NT control sgRNAs by one-way ANOVA with Dunnett’s post-test. **e**, CRISPRi effects on intracellular MUC5AC were quantified by flow cytometry. **f**, Targeting SPDEFe but not surrounding regions inhibits IL-13-induced MUC5AC production. sgRNAs used in (**b**-**f**) were compared with sgRNAs targeting flanking regions ∼2 and 4 kb away from SPDEFe (magenta) in a separate set of experiments (three donors, two replicates per donor). MUC5AC-producing cells relative to the mean of MUC5AC-producing cells from IL-13-stimulated cells with NT gRNAs in the same donor are displayed in percentage. ns, not significant; **, *p* < 0.01; and ****, *p* < 0.0001 compared with IL-13-stimulated HBECs with NT-1 sgRNA by one-way ANOVA with Tukey’s post-test. A full table of comparisons is in Supplementary Table 1. **g**-**j**, CRISPRi effects on mucin staining and mucociliary transport. HBECs were treated as above using either NT-2 gRNA, a SPDEF-TSS(+34) gRNA, or a SPDEFe(+87) gRNA. Sections (**g**) and extracellular mucus gels from whole mount preparations (**h**) were stained for MUC5AC (cyan), MUC5B (red), the ciliated cell marker ac-α-Tub (yellow). Nuclei were stained with DAPI (purple). Scale bars: 20 *μ*m (**g**) and 100 *μ*m (**h**). Images are representative of two experiments with different donors. **i** and **j**, Mucociliary transport rates were determined from trajectories of fluorescent microspheres placed on gels atop cells. **i**, Superimposition of 10 images acquired at 1-s intervals. Scale bars: 50 *μ*m. **j**, Microsphere speeds determined from three donors, one well per donor, four fields per well (total of 12 fields per condition). Values represent median microsphere speed for each field. Boxes extend from the 25th to the 75th percentile, the horizontal line within the box indicates the mean, and the whiskers represent minimum and maximum values. *, *p* < 0.05 and **, *p* < 0.01 by one-way ANOVA with Tukey’s post-test.

We next analyzed whether targeting SPDEFe could reverse IL-13-induced abnormalities in mucus organization and function that can contribute to asthma morbidity and mortality. We previously reported that IL-13 stimulation induces the formation of MUC5AC-rich mucus that remains tethered to the epithelium, resulting in impaired mucociliary transport^21^. Consistent with our analyses of the effects of SPDEFe gRNAs on *MUC5AC* mRNA and MUC5AC protein by flow cytometry, we found that targeting SPDEFe prevented the formation of MUC5AC-rich mucus domains (Fig. 3g and h). We investigated whether targeting SPDEFe via CRISPRi could also rescue mucociliary transport abnormalities induced by IL-13 (Fig. 3i and j). IL-13 stimulation of cells expressing a non-targeting sgRNA led to a marked reduction in mucociliary clearance. Targeting the *SPDEF* TSS or SPDEFe completely rescued the cell cultures from IL-13-induced mucostasis. These results show that direct targeting of SPDEFe prevents IL-13 induction of *SPDEF*, the goblet cell transcriptional program, and impaired mucociliary clearance.

### The *SPDEF* enhancer is IL-13-inducible and cell type-selective

To better understand the relationship between SPDEFe and IL-13 responsiveness, we directly tested the activity of SPDEFe in a GFP-based reporter assay^22^. A 589-bp sequence from SPDEFe drove reporter expression in an IL-13-dependent manner in a subset of HBECs cultured and differentiated at ALI (Fig. 4a). To characterize this subset, we dissociated differentiated HBECs and analyzed reporter expression by flow cytometry using a panel of cell type markers^23^. The activity of SPDEFe was significantly induced by IL-13 only in secretory cells (Fig. 4b-e). To determine whether this reflected a cell type-specific difference in the IL-13/STAT6 signaling pathway, we tested a reporter construct containing a germline Ige promoter (IGHEe)^24^. IGHEe contains CEBPβ and STAT6-binding sites and has a known role in IL-4-induced STAT6-mediated B cell immunoglobulin class switching. IL-13 induced IGHEe activity in all cell types, demonstrating that the cell type selectivity of SPDEFe is not attributable to difference in activation of the STAT6 pathway. To further assess the dependence on cell differentiation, we transduced HBECs that were maintained in standard (submerged) cell culture, where cells remain in a poorly differentiated state, and two lung epithelial cell lines, BEAS-2B and A549 (Fig. 4f). IL-13 induced the IGHEe reporter but not the SPDEFe reporter in these cells. These findings indicate that SPDEFe is an IL-13-dependent enhancer that is selectively active in differentiated secretory cells.

**Figure 4.**
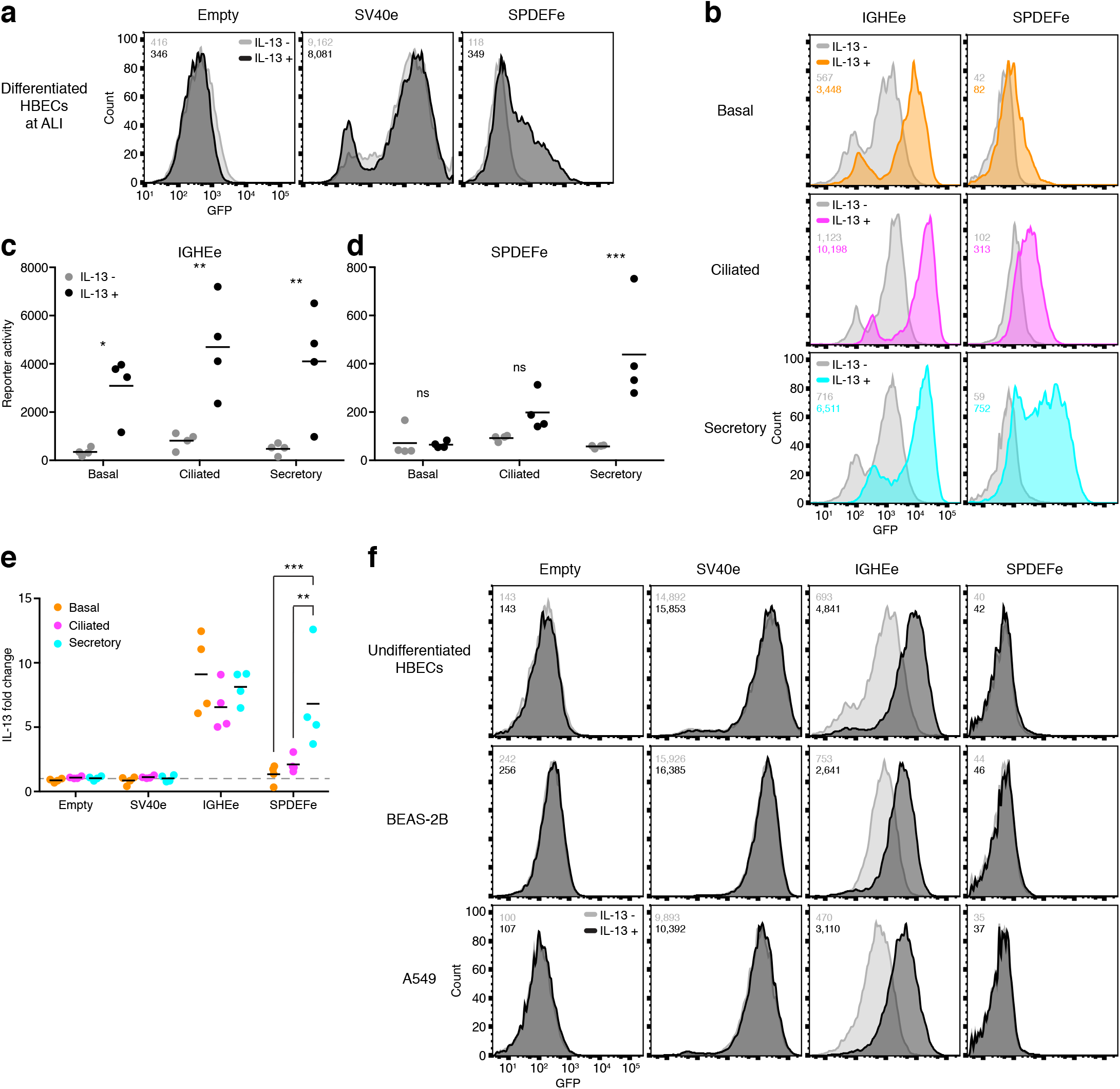
The *SPDEF* enhancer drives IL-13-inducible cell type-selective transcription. **a**, HBECs were transduced with lentiviral GFP reporter constructs of containing only a minimal promoter (Empty), a positive control SV40 enhancer (SV40e), and SPDEFe (chr6:34,586,344-34,586,932), differentiated at ALI without (grey) or with (black) IL-13 stimulation during the last 7 d of culture. Representative histograms from one of at least three donors are shown, and mean fluorescence intensities are displayed. **b**-**e**, IGHEe- and SPDEFe-driven reporter expression in basal (NGFR^+^ CEACAM6^-^), ciliated (a-Ac-Tub^+^), and secretory (CEACAM6^+^ NGFR^-^) cells. **b**, Representative histograms from one out of four donors with mean fluorescence intensities. **c**-**d**, Mean fluorescence activity for all donors. ns, not significant; *, *p* < 0.05; and **, *p* < 0.01 for comparison between with and without IL-13 stimulation by two-way ANOVA with Sidak’s post-test. **e**, IL-13-induced fold changes in reporter expression in each cell type. **, *p* < 0.01 and ***, *p* < 0.001 for comparison between cell types by two-way ANOVA Tukey’s post-test; all other differences were not statistically significant. **f**, Reporter activity in undifferentiated HBECs and two lung epithelial cell lines, BEAS-2B and A549. Representative histograms from at least three donors or experiments are shown, and mean fluorescence intensities are displayed.

Although overproduction of MUC5AC is detrimental in asthma, this mucin has been shown to protect against influenza infection in viral models. Whereas there is typically minimal *MUC5AC* expression in unstimulated HBECs under our usual ALI culture conditions, an alternative cell culture medium, PneumaCult-ALI, supports the differentiation of an epithelium that more closely models that seen *in vivo* in that it contains MUC5AC-producing cells that develop without a requirement for IL-13 stimulation^25^. When cultured using PneumaCult-ALI, HBECs produced MUC5AC but did not activate the SPDEFe reporter (Supplementary Fig. 5a-c). *MUC5AC* expression is also induced by IL-1b in the context of airway infection^26^. We found that IL-1b induction of *MUC5AC* was not accompanied by activation of the SPDEFe reporter (Supplementary Fig. 5d-f). These results demonstrate that the SPDEFe activation is regulated by IL-13 but not by the two other *MUC5AC-*inducing stimuli that we studied.

### SPDEFe activity is mediated by STAT6 and KLF5

To further characterize SPDEFe and define the minimal active region, we tested overlapping ∼150 bp fragments spanning the 589-bp SPDEFe sequence. Of the seven constructs tested, only the first two most proximal were active in IL-13-stimulated HBECs (Fig. 5a). The first fragment, SPDEFe(1-147), encompassing the first 147 bp of the sequence and overlapping with the second fragment, gave the strongest response and was sufficient for cytokine inducibility and secretory cell selectivity (Fig. 5b). SPDEFe(1-147) includes predicted binding sites for several transcription factors and DNA-binding proteins including STAT6, which has a known role in IL-13 signaling. Among those, only *KLF5* was significantly more abundant in secretory cells compared to basal and ciliated cells by scRNA-seq (1.4- and 1.5-fold higher, respectively; FDR = 1.3 × 10^−91^ and 6.1 × 10^−51^). We mutated the SPDEFe(1-147) STAT6 and KLF5 predicted binding sites and tested the effects using the lentiviral reporter assay (Fig. 5c). Mutating any of the two STAT6 or two KLF5 binding sites reduced or eliminated IL-13-induced enhancer activity. These findings demonstrate the requirement for STAT6 and KLF5 predicted binding sites within a 60-bp (44-103) region of SPDEFe.

**Figure 5.**
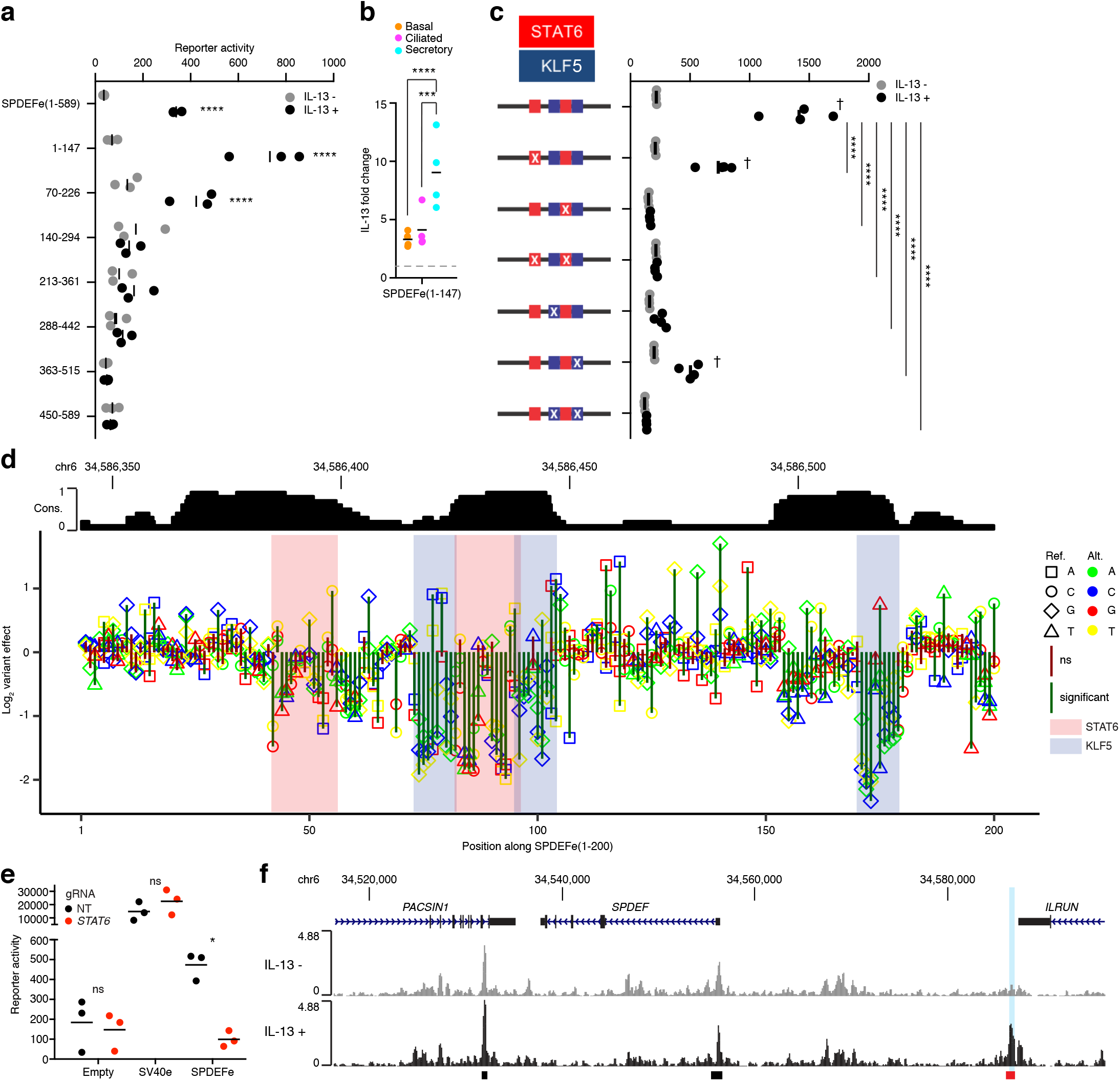
SPDEFe activity requires STAT6 and KLF5 binding sites. **a**, Seven overlapping ∼150-bp fragments of 589-bp SPDEFe were tested via the lentiviral reporter assay in HBECs cultured at ALI without (grey) or with (black) IL-13 stimulation. ****, *p* < 0.0001 for comparison between with and without IL-13 stimulation by two-way ANOVA with Sidak’s post-test (three donors, one well per donor). **b**, IL-13-induced activity of the SPDEFe(1-147) fragment in each cell type. These data were acquired as a part of the same experiment performed in Fig. 4e. ***, *p* < 0.001 and ****, *p* < 0.0001 for comparison between cell types by two-way ANOVA with Tukey’s post-test. **c**, Mutations were made in core sequences of predicted STAT6 and KLF5 binding sites (JASPR) in SPDEFe(1-147) and tested via the lentiviral GFP reporter assay in 3 donors. †, *p* < 0.0001 for comparison between with and without IL-13 stimulation and ****, *p* < 0.0001 for comparison against the reference SPDEFe(1-147) sequence by two-way ANOVA with Sidak’s post-test. All other differences were not statistically significant (a-c). **d**, Saturation mutagenesis MPRA of bases 1-200 of SPDEFe. Values represent log_2_ effects of each single nucleotide substitution (combined data from two donors). Positions of predicted STAT6 and KLF5 binding sites (JASPR) are highlighted in red and blue, respectively. The extent of sequence conservation within 30 mammalian species by PhastCons is displayed at the top. **e**, Effects of *STAT6* targeting on reporter expression. HBECs were transfected with rCas9 and either NT-2 control (black) or *STAT6* (red) sgRNA prior to introduction of the reporter constructs. ns, not significant and *, *p* < 0.05 for comparison of NT-2 and *STAT6* gRNAs by *t*-test with multiple comparisons correction using the Holm-Sidak method. **f**, CUT&Tag analysis of KLF5 binding near the *SPDEF* locus. Tracks represent mean reads per million from six experiments (three donors, two replicates per donor) involving HBECs cultured without (grey) or with (black) IL-13 stimulation. Called peaks are indicated by bars below the tracks; the sole IL-13-enriched peak (FDR < 0.1) is in red. SPDEFe is highlighted in cyan.

We used a massively parallel reporter assay (MPRA)^27,28^ for finer mapping of the enhancer. All 600 possible single nucleotide variants of the first 200 bp of SPDEFe (SPDEFe(1-200)) were compared with the reference sequence (Supplementary Data 4). MPRA measurements from HBECs from two donors were well correlated (*R =* 0.77, *p* < 2.2 × 10^−16^, Supplementary Fig. 6a) and were pooled for further analysis. Of the 600 mutations tested, 31 increased reporter expression and 128 reduced expression in IL-13-stimulated cells (absolute fold-change ≥ 1.5, FDR < 0.05) (Fig. 5e). Function-perturbing mutations clustered at the previously identified STAT6 and KLF5 motifs and at a third KLF5 motif (positions 170-179), all of which were well conserved in mammals. Effects of mutations in the sequences with STAT6 motifs tended to be larger when they involved core positions in the 5’-TTCNNNNGAA-3’ motif (Supplementary Fig. 6b and c). For sequences containing the KLF5 motif, mutations that reduced similarity to the consensus motif tended to reduce activity, whereas some mutations that increased similarity to the consensus increased activity (Supplementary Fig. 6d-f). 27 mutations that we tested correspond to SNPs found in NCBI dbSNP release 153; all of these are rare (frequency < 1%). Of those, 2 increased and 6 decreased enhancer activity (FDR < 0.05). The SNP rs1165834930 G/C in the third KLF5 binding site led to the largest effect among all variants (5.0-fold). These results demonstrate that sequences containing STAT6 and KLF5 binding sites are required for SPDEFe activity and that sequence variants found in human populations alter the activity of this regulatory element.

We complemented studies of the *cis*-regulatory elements with studies of STAT6 and KLF5 *trans*-factors. CRISPR-mediated deletion of STAT6 prevented IL-13-induced activation of the SPDEFe reporter (Fig. 5f). Taken together with the effects of mutating the STAT6 motifs, this suggests a direct role for STAT6 in SPDEFe activation. CRISPR targeting of *KLF5* in HBECs led to cell death upon culturing at ALI, consistent with a previous report that *Klf5* is required for airway epithelial maturation in mice^29^. To assess whether KLF5 binds to SPDEFe, we used Cleavage Under Targets and Tagmentation (CUT&Tag)^30^ to identify KLF5-associated genomic regions in HBECs. The KLF5 motif was highly enriched in 22,948 regions identified using CUT&Tag (*p* = 10^−1306^). 863 regions were enriched by IL-13. Of three KLF5-associated regions detected within 50-kb of the *SPDEF* TSS, one was enriched by IL-13, and this region contained SPDEFe (Fig. 5g). This result indicates that IL-13-induced activation of SPDEFe is associated with increased binding of KLF5 to this enhancer.

### SPDEFe can be repurposed to target pathologic mucus production

To test whether a cell type-dependent, cytokine-activated regulatory element could be used as a therapeutic tool for selectively modulating expression of genes that contribute to disease, we used SPDEFe to drive CRISPRi of either *SPDEF* or *MUC5AC*. SPDEFe was placed upstream of a minimal promoter controlling the expression of *KRAB*-*dCas9* in a lentiviral vector. SPDEFe-*KRAB-dCas9* lentiviruses were transduced into HBECs together with lentiviruses encoding gRNAs targeting the *SPDEF* or *MUC5AC* TSS (Fig. 6a). SPDEFe drove expression of *KRAB-dCas9* in an IL-13-inducible manner (Fig. 6b). Substitution of SPDEFe with sequences containing two or three repeats of the first 147 bp of SPDEFe increased IL-13-driven *KRAB-dCas9*. Induction of *KRAB-dCas9* in cells expressing the *SPDEF* gRNA significantly reduced *SPDEF* and *MUC5AC* mRNA levels (Fig. 6c and d) as well as MUC5AC protein level Fig. 6e). CRISPRi of *MUC5AC* showed significant IL-13-induced *MUC5AC* mRNA and protein reduction without effects on *SPDEF* expression. To verify that the ability of the *SPDEF* enhancer sequences to suppress *MUC5AC* expression is IL-13-specific, we tested the effect of our regulatory circuit on IL-1β-induced *SPDEF* and *MUC5AC* expression. IL-1β did not induce the expression of *KRAB-dCas9* (Fig. 6f) and SPDEFe-based CRISPRi did not prevent IL-1β induction of *SPDEF* and *MUC5AC* (Fig. 6g and h). We conclude that regulatory circuits based on use of SPDEFe to regulate CRISPRi can be used to selectively inhibit IL-13-induced *MUC5AC* expression.

**Figure 6.**
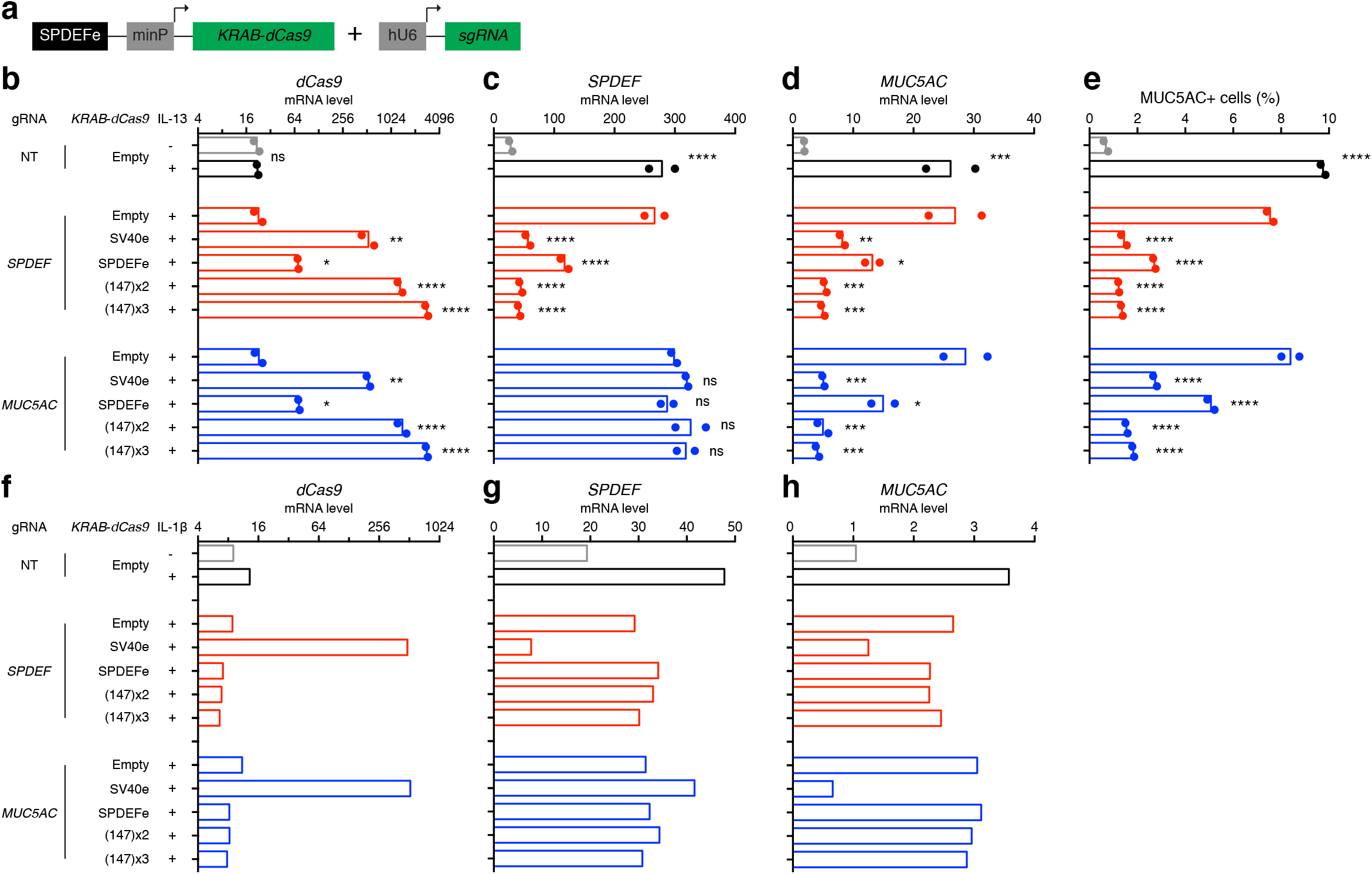
SPDEFe can be used to prevent IL-13-induced mucin production by secretory cells. **a**, A lentivirus containing a *KRAB*-*dCas9* transgene driven by SPDEFe (or other enhancers) and a second lentivirus driving expression of sgRNA targeting *SPDEF* (SPDEF-TSS(+34)) or *MUC5AC* (MUC5AC-TSS(+134)) promoter were used in combination. minP, minimal promoter. Transduced cells were cultured at ALI without cytokine stimulation or with IL-13 (**b**-**e**) or IL-1b (**f**-**h**) stimulation for the last 7 days of culture, as indicated. Changes in expression of *dCas9* (**b** and **f**), *SPDEF* (**c** and **g**), and *MUC5AC* (**d** and **h**) were measured by qRT-PCR. MUC5AC-producing cells (**e**) were quantitated by flow cytometry. IL-13 data (**b**-**e**) are from two experiments with different donors. *, *p* < 0.05; **, *p* < 0.01; ***, *p* < 0.001; and ****, *p* < 0.0001 compared with unstimulated Empty vector control for NT gRNA and IL-13-stimulated Empty vector control for *SPDEF* and *MUC5AC* gRNAs by one-way ANOVA Tukey’s post-test. The IL-1β data (**f**-**h**) are from a single experiment with one donor, and no statistical analysis was performed.

## Discussion

Pleiomorphic effects of cytokines on airway epithelial cells play a central role in asthma. The airway epithelium comprises multiple cell types, but the effects of cytokines on specific cell types and the mechanisms underlying cell type-selective responses are incompletely understood. We performed genome-wide transcriptomic analyses by bulk and single cell RNA sequencing in HBECs in response to the asthma-associated cytokines IFN-α, IL-17, and IL-13. We found that though all tested cytokines induced transcriptional responses, more cell type-selective response was observed in different airway epithelial subsets upon IL-13 stimulation. Epigenomic analysis showed that IL-13 effects were associated with changes in H3K27ac, a marker of active promoters and enhancers, near IL-13-regulated genes. To obtain a detailed understanding of how IL-13 induces cell type-specific responses in the airway epithelium, we focused on the regulation of *SPDEF*, an IL-13-inducible gene that is required for pathologic mucus production. We identified an IL-13-activated distal regulatory element located ∼30 kb from the *SPDEF* TSS. CRISPRi targeting of this element inhibited *SPDEF* expression, and the element was sufficient to drive IL-13-regulated cell type-specific expression of a reporter transgene. Based on these findings, we designed and implemented a synthetic regulatory circuit that selectively inhibits IL-13-induced pathologic mucus production and mucostasis. This work demonstrates a critical role for enhancers in driving cell type-specific cytokine responses and illustrates how these enhancers could be exploited for new therapeutic approaches.

The cytokines that we studied affect different subsets of individuals with asthma^3,5,6^. We found that IL-13, IL-17, and IFNs have highly distinct effects on HBECs. IFN stimulation of HBECs activated many IFN-stimulated pathways previously identified in other cell types, including those involved in anti-viral immunity and inflammatory responses. The effects of IFN-α stimulation were remarkably similar in each of the three major HBEC subsets. IL-17 induced genes involved in anti-microbial immune responses. In comparison to IFN-α, more IL-17 effects were cell type-selective. IL-13 had the most prominent cell type-selectivity. A previous report focusing on IL-13 effects on human tracheal epithelial cells also showed substantial cell type-specific effects^31^. In basal cells, IL-13 selectively induced genes encoding eotaxin-3 (CCL26), which recruits eosinophils^32^, and POSTN, a pro-fibrotic and anti-inflammatory matricellular protein^33^ that has been used as a marker for type 2-high asthma^34^. *SERPINB2*, another type 2-high asthma marker, was among the genes most highly induced in ciliated cells. A substantial set of genes were selectively induced by IL-13 in secretory cells. Many of these genes have been associated with goblet cell metaplasia, a process that transforms the histologic appearance and secretome of secretory cells. Goblet cells are the source of tethered MUC5AC, which impairs mucociliary clearance and causes airway obstruction^21^. Hence, IL-13 has distinct effects even within epithelial cells from a single tissue, and these cell type-selective effects contribute to different features of asthma.

In seeking a mechanism to account for cell type-selective effects of IL-13 on gene expression, we focused on activation of DNA regulatory elements. IL-13 induced a substantial number of changes in H3K27ac. Furthermore, IL-13-induced increases in H3K27ac were associated with increased expression of nearby genes, suggesting a functional role. In contrast, we found that the profound effects of IFN-α or IL-17 stimulation on HBEC gene expression were not accompanied by substantial changes in H3K27ac. Type I interferon signaling has been shown to drive chromatin remodeling and post-translational histone modifications in prior studies, and it is possible that the lack of H3K27ac changes after IFN-α stimulation in our experiments relates to the relatively shorter duration of cytokine stimulation (1 d for IFN-α versus 7 d for IL-13).

To gain a detailed understanding of how enhancers can contribute to cell type-selective responses, we focused on a novel enhancer located 30 kb from the TSS of *SPDEF*, a transcription factor that is critical for goblet cell metaplasia. Deletion of *Spdef/SPDEF* in mice^19^ or HBECs^20^ prevented IL-13-dependent induction of goblet cell metaplasia. To determine whether the SPDEFe candidate enhancer identified by ChIP-seq regulates *SPDEF*, we made use of a CRISPRi-based approach for discovering regulatory elements and identifying their target genes^37,38^. Targeting the enhancer using CRISPRi led to reduced expression of *SPDEF* and prevented SPDEF-dependent goblet cell metaplasia, pathological mucus production, and mucostasis. This effect was localized and specific. Sequences from the enhancer conferred the ability to transcribe a reporter gene in an IL-13-dependent manner. Remarkably, this activity was selective for secretory versus basal or ciliated cells and was completely absent in undifferentiated epithelial cells. This specificity could not be explained by a global difference in IL-13 signaling, since a STAT6-binding enhancer previously identified in B lymphocytes conferred robust IL-13-responsive transcription in all cell types that we studied.

We used complementary approaches to dissect the *cis*-regulatory sequences and associated DNA-binding *trans*-factors required for selective IL-13-dependent transcription in secretory cells. Mutagenesis of two STAT6 motifs and three KLF5 motifs reduced enhancer activity. The enhancer was inactive in STAT6-deficient cells, suggesting that IL-13 activation of the STAT6 signaling pathway leads directly to enhancer activation, although we cannot formally exclude an indirect role for STAT6. Mutations that disrupted the KLF5 motifs reduced activity whereas some mutations that produced sequences closer to a known KLF5 consensus motif increased activity. Our finding that KLF5 was physically associated with enhancer region in an IL-13-dependent manner in combination with our mutagenesis results strongly suggest KLF5 has a direct role. Reported mechanisms of target gene activation by KLF5 include interaction with TFIIB and TATA box-binding protein, other transcriptional coregulators, and post-translational modifiers including HDAC1/2 and p300^39^. Our results are consistent with a model where IL-13 stimulation triggers STAT6 activation, recruitment of both STAT6 and KLF5, and subsequent KLF5-mediated H3K27ac, but our studies were performed late (7 d) after addition of IL-13, and further studies will be required to determine the sequence of events. *KLF5* expression and function may help account for differences in enhancer activity between airway epithelial cell types, since we found modestly higher levels of *KLF5* mRNA in secretory cells compared with basal and ciliated cells. It is also plausible that other secretory cell-specific *trans*-factors could play a role by interacting with KLF5 or STAT6. Previous reports of IL-4 effects on macrophages found functional interaction between STAT6 and another Kruppel-like factor, KLF4, at the *Arg1* promoter^40^ and enrichment of STAT6 and KLF motifs in IL-4-induced enhancers^41^, suggesting that direct or indirect interactions between STAT6 and KLF family members may be of more general importance in IL-4 and IL-13 signaling.

Our discovery of an enhancer that is sufficient for a cell type-selective IL-13 response led us to test whether this could be used to engineer a synthetic regulatory circuit to control pathologic responses to IL-13 in the airway epithelium. CRISPR-based approaches for activating or interfering with gene expression provide useful experimental tools and hold great promise for therapy. For example, a CRISPR-based activation (CRISPRa) system corrected obesity caused by haploinsufficiency of *Sim1* in mice^42^. In many cases, it is important to limit gene activation or repression to particular cell types or conditions. In the case of goblet cells and mucus production, suppressing IL-13-induced goblet cell metaplasia and *MUC5AC* overexpression would likely be beneficial for preventing airway obstruction in asthma, but global suppression of goblet cells and *SPDEF* and *MUC5AC* expression might be deleterious for host defense. Various approaches including cell type-specific CRISPR/Cas9 delivery vehicles^43^, transcript-specific riboswitches^44^, and miRNA-responsive switches^45^ can be useful for targeting CRISPR-based systems to specific cell types. As an alternative, we used a simple two-component system comprised of a cell-type selective enhancer to control production of KRAB-dCas9 together with a sgRNA directed against the *SPDEF* promoter. Using this system, we were able to efficiently suppress IL-13-induced goblet cell production. Using a different gRNA against *MUC5AC* promoter allowed us to target this critical downstream gene without disrupting *SPDEF* expression. By using an enhancer that is not activated by other stimuli that promote *SPDEF* and *MUC5AC* expression, including PneumaCult-ALI medium and IL-1β, our approach selectively suppresses only IL-13-driven pathology. These results demonstrate that incorporating cell type-selective cytokine-responsive enhancers into synthetic regulatory circuits could provide a simple but powerful tool for highly selective treatment of human diseases. Identifying additional enhancers that drive cell type-selective responses to IL-13 and other cytokines would broaden our understanding of how cytokines exert their pleiotropic effects and provide more tools for future therapies.

## Methods

All methods are available in Supplementary Information.

## Author contributions

K.D.K., L.R.B., W.L.E., N.A., and D.J.E. contributed to study conception and design. K.D.K., L.R.B., O.Y.B., X.Z., D.I.S., and L.T.Z. performed experiments. K.D.K., L.R.B., W.L.E., J.S., O.Y.B., and X.Z. analyzed data. K.D.K., L.R.B., W.L.E., and D.J.E. interpreted data. W.E.F., N.A., and D.J.E. supervised study execution. K.D.K. and L.R.B. drafted the manuscript. K.D.K., L.R.B., W.L.E., J.S., O.Y.B., X.Z., D.I.S., L.T.Z., W.E.F., N.A., and D.J.E. revised the manuscript.

## Acknowledgments

We thank Michael Matthay and Paul Wolters (UCSF) for supplying human bronchial specimens and Jane Gordon, Sarah Elmes (Laboratory for Cell Analysis at UCSF), Kari Herrington (Nikon Imaging Center at UCSF), Eric Chow (Center for Advanced Technology at UCSF), Siranoosh Ashtari (DNA Technologies & Expression Analysis Core Laboratory at UC Davis), Luke Gilbert (UCSF), and Jonathan Weissman (MIT) for technical advice and assistance. This work was supported in part by the National Institute of Health grants U19 AI077439 (D.J.E.), R35 HL145235 (D.J.E. and N.A.), R01 HL117004 (N.A.), K99/R00 HL135404 (W.L.E.); a Sandler Foundation Program for Breakthrough Biomedical Research Postdoctoral Independent Research Award 7028238 (L.R.B.); and a National Natural Science Foundation of China grant 81970016 (X.Z.).

## Notes

### Competing Interest Statement

The authors have declared no competing interest.

