## Supplementary Information for "Targeting a central feature of asthma using a cell type-selective IL-13-responsive enhancer"

†Equal contributions

*Corresponding

**Supplementary Materials & Methods**

**Cells**

The UCSF Committee on Human Research approved the use of HBECs isolated from lung transplant recipients' explanted lungs or lungs not used for transplantation. Written consent was not required since materials were leftover clinical samples obtained from de-identified individuals. HBECs were seeded onto 10-cm dishes coated with human placental collagen (HPC; Sigma-Aldrich, St. Louis, MO) and propagated in BEGM (Lonza, Walkersville, MD) with 10 $\mu$M Y-27632 (Enzo Life Sciences, Farmingdale, NY) as described previously[1,2]. BEAS-2B and A549 cells were cultured in 1:1 DMEM/F12 medium containing Non-Essential Amino Acids (Genesee Scientific, San Diego, CA) and DMEM, respectively, with 10% FBS (Thermo Fisher Scientific, Waltham, MA).

**Plasmids**

Plasmid hU6-sgRNA-cr3-EF1a-Puro was made by cloning in hU6-sgRNA-cr3-EF1a-Puro construct from pMJ117[3] (a gift from J. Weissman lab at MIT) into XhoI,XbaI-cut (New England BioLabs, Ipswich, MA) backbone of BTV[4]. Plasmid EF1a-dCas9-KRAB-P2A-BSD was made by cloning in EF1a-dCas9, KRAB from IGI-P0152 (a gift from J. Corn lab at ETH Zurich via L. Gilbert lab at UCSF), P2A-BSD from pLenti-SFFV-KRAB-dCas9-P2A-BSD into XhoI,XbaI-cut backbone of BTV. Plasmid mP-KRAB-dCas9_EF1a-BSD was made by cloning in SV40 poly(A) signal from BTV, BSD, EF1a from lentiCRISPRv2, antirepressor#40-mP from pLS-mP[5], and KRAB-dCas9 from pHR-

TRE3G-KRAB-dCas9-P2A-mCherry[6] (a gift from L. Gilbert lab) into XhoI,XbaI-cut backbone of BTV. NEBuilder HiFi DNA Assembly (New England BioLabs) was used for cloning.

**Lentivirus preparation**

Third generation lentivirus was produced by transfecting appropriate plasmids or plasmid libraries with three lentivirus-packaging plasmids (pMDL, pVSV-G, and pRSV-Rev) into HEK293T cells using TransIT-293 Transfection Reagent (Mirus Bio, Madison, WI). Harvested lentivirus was concentrated 100-fold using Lenti-X Concentrator (Takara Bio, Mountain View, CA), resuspended in PBS, and stored at -80 °C in aliquots.

**Bulk RNA-seq**

HBECs from six donors (1 = 12-43, 2 = 13-26, 3 = 13-33, 4 = 13-24, 5 = 13-23, and 6 = 13-28) were cultured at ALI for 23 d, as described previously[2,7], without cytokine or with IFN-$\alpha$ (10 ng/mL for the final 1 d), IFN-$\gamma$ (10 ng/mL for the final 1 d), IL-13 (10 ng/mL for the final 7 d), IL-17 (10 ng/mL for the final 7 d), or the combination of IFN-$\alpha$ and IL-13. IFNs and ILs were acquired from R&D Systems (Minneapolis, MN) and Peprotech (Rocky Hill, NJ), respectively. After removal of accumulated mucus with 10 mM dithiothreitol (DTT; Enzo Life Sciences, Farmingdale, NY) in PBS, cells were trypsinized with 0.25% trypsin-EDTA (Thermo Fisher Scientific) for 10-15 min at 37 °C and neutralized. RNA was isolated from harvested HBECs, and bulk RNA-seq was performed as previously described[8,9]. We have previously reported other analyses based upon a subset of these data from unstimulated cells and cells stimulated with IFN-α[8] and IL-17[9]. In addition, data for 332 genes associated with SARS-CoV-2 have been reported in another study[10]. Raw read counts per sample and gene were used as input to DESeq2 for unsupervised and supervised analysis. Read counts were first normalized for sequencing depth and transformed using DESeq2's variance stabilizing transformation. Donor specific impacts on the stabilized counts where then removed with LIMMA's removeBatchEffect function. The major sources of variability where then inspected with a principal component analysis (PCA) using the top 3000 most variable genes across the experiment. Differential gene expression in response to each cytokine treatment compared to untreated was then determined using DESeq2 after correction for the donor specific effects using the design formula: ~ donor + condition, where condition includes all treatment options, IFN-α, IFN-γ, IL-13, IL-17, and the combination of IFN-α and IL-13. Pairwise comparisons between treatments and the untreated control were carried out with DESeq2 results function using a Wald test to estimate *p*-value and a multiple testing correction using the FDR method.

**Single cell RNA-seq**

We used HBECs from four (1 = 12-43, 2 = 13-26, 3 = 13-33, and 6 = 13-28) of the six donors used for bulk RNA-seq. Cells were cultured at ALI for 23 d without cytokine or with IFN-α (10 ng/mL for the final 1 d), IL-13 (10 ng/mL for the final 7 d), IL-17 (10 ng/mL for the final 7 d), or the combination of IFN-α and IL-13. Single cell suspensions were generated as described above. Cells were manually counted using a hemocytometer, and equal numbers of cells from each donor were pooled. Each pool included four samples (one from each donor and each representing a different cytokine stimulation). 10x Genomics scRNA-seq libraries (Pleasanton, CA) were prepared[11] and sequenced using Illumina NovaSeq 6000 S4 flow cells (San Diego, CA) at the Center for Advanced Technology at UCSF. Resulting sequence reads were processed into a cell-by-gene counts matrix using 10x Genomics Cell Ranger software. Single nucleotide polymorphisms that were unique to one of the four donors were identified from the bulk RNA-seq data following the GATK best practices[12] and used to assign each cell to the appropriate donor using Demuxlet[13]. Demultiplexed droplets uniquely assigned to a single individual were then processed using Seurat[14,15] with the following filters: nFeature_RNA >= 500 & nFeature_RNA < 7500 & nCount_RNA > 1200 & percent.mito < 0.5 & percent.ribo < 0.35. The top 10,000 variable genes were carried forward for normalization, unsupervised clustering, UMAP visualization, and cell type identification. Normalized data was scaled while regressing out the 10x Genomics well effect, percent mitochondrial reads, percent ribosomal reads, and the number of unique molecular indexes per cell. Cells from cultures that were not stimulated with cytokines were then isolated for clustering and cell type identification. These unstimulated cells then served as the reference dataset for Seurat's reference-based multiple dataset integration using SCTransform[16] with label transfer in order to use anchor genes from the unstimulated cells to identify cell types across all samples. Cell identities were then returned to the original Seurat object prior to SCTransform for further investigation. We used Seurat's FindAllMarkers to identify cell type marker genes. To find genes that have cell-type-specific responses to cytokine stimulation, we used the package MAST[17]. A hurdle model was fitted with the covariates batch, individual and cngeneson (a MAST-specific covariate that corrects for cellular detection rate), cell type, and cell type by cytokine interaction terms. Significance testing using the lrt() function was used to find differentially expressed genes by cytokine stimulation as compared to untreated genes in the same cell type. Data for 332 genes associated with SARS-CoV-2 infection have been reported in another study[10]. For linear model cell type interaction analysis, we downsampled the number of cells in basal, ciliated, and secretory cells to 450 cells each, to minimize the effect of sample size on power of detecting differentially expressed genes. The same hurdle model was fitted. Genes with *p*-value for cell type by cytokine interaction terms < 0.1 were called significant. Gene set enrichment analysis was performed using the package iDEA[18] and Gene Ontology Biological Process (GOBP) gene sets from Msigdb ("c5.go.bp.v7.4.symbols.gmt"). We used the default parameters from the function iDEA.fit, except that the minimum number of DE genes was set to 2. $Log_2$ fold change and *p*-values from MAST-based differential gene expression were used as input.

**H3K27ac ChIP-seq**

H3K27ac ChIP-seq was performed on three of the donors used in sequencing studies (1 = 12-43, 2 = 13-26, and 6 = 13-28) and one additional donor 16-05. HBECs were cultured at ALI for 23 d without cytokine or with IFN-$\alpha$ (10 ng/mL for the final 1 d), IL-13 (10 ng/mL for the final 7 d), or IL-17 (10 ng/mL for the final 7 d). ChIP-seq was performed using previously reported methods[19]. In brief, cells were harvested, fixed with 0.5% paraformaldehyde (PFA; Thermo Fisher Scientific), and lysed, and then chromatin was sheared using a Covaris S2 sonicator (Woburn, MA). Chromatin immunoprecipitation was performed with an H3K27ac antibody (ab4729, rabbit IgG; Abcam, Cambridge, UK) using the Diagenode LowCell# ChIP kit (Denville, NJ) as per the manufacturer's instructions. Libraries were prepared using the Rubicon DNA-seq kit (Rubicon Genomics, Atlanta, GA) and then sequenced on an Illumina HiSeq 4000 using 50-bp single end at the Center for Advanced Technology at UCSF. Sequencing reads were aligned by STAR v2.5.2b[20], and reproducible peaks were called for each condition using the ENCODE IDR framework[21]. Additionally, regions blacklisted by the ENCODE consortium[22] for their tendency to produce anomalous results were removed from the peak set using bedtools. Analysis for differentially enriched peaks in response to cytokines was carried out as previously described[23]. Differentially modulated peaks were inspected for proximity to differentially expressed genes from bulk and single cell RNA-seq data using Bedtools v2.29.2[24]. HOMER motif analysis (http://homer.ucsd.edu/homer/introduction/programs.html)[25] was performed on upregulated and downregulated H3K27ac peaks with parameter size –given. Peaks with *p*-value > 0.1 was used as background peaks.

**CRISPRi targeting of genomic regulatory elements**

Each gRNA sequence (Supplementary Table 2) was selected using CRISPick (https://portals.broadinstitute.org/gppx/crispick/public) and GuideScan[26] and was used to design a pair of complementary 5′-phosphorylated oligonucleotides (5′-$PO_4$-ATGNNNNNNNNNNNNNNNNNNNNGTTTCAGAGC-3′ and 5′-$PO_4$-TTAGCTCTGAAACNNNNNNNNNNNNNNNNNNNNCATGTTT-3′; where Ns indicate sequences that vary according to the gRNA). A G was added to the 5′ ends of gRNA sequences not beginning with a G. gRNA oligonucleotides (oligos; IDT, San Jose, CA) were annealed and cloned into the hU6-sgRNA-cr3-EF1a-Puro plasmid digested with BstXI and BlpI (New England Biolabs). HBECs were initially transduced with EF1a-dCas9-KRAB-P2A-BSD lentivirus and maintained in BEGM with 10 μM Y-27632 and 10 μg/mL blasticidin (Thermo Fisher Scientific) for 3 d for selection. Cells were subsequently transduced with sgRNA lentivirus and maintained in BEGM with 10 μM Y-27632, 10 μg/mL blasticidin, and 1 μg/mL puromycin (Thermo Fisher Scientific) for 3 d for selection. Transduced and selected cells were then passaged to Transwell inserts for culture at ALI for 23 d. Where indicated, IL-13 (10 ng/mL) was added for the final 7 d of culture.

**Quantification of mRNA transcripts**

mRNAs were measured by quantitative real-time RT-PCR (qRT-PCR). RNA was extracted from HBECs using the RNeasy Mini Kit (QIAGEN, Hilden, Germany) or the RNA/DNA/Protein Purification Plus Kit (Norgen Biotek, Thorold, ON, Canada) according to manufacturers' instructions. RNA was reverse-transcribed using SuperScript III First-Strand Synthesis System (Thermo Fisher Scientific). cDNA was analyzed by qRT-PCR (PowerUp SYBR Green, Thermo Fisher Scientific; primer sequences in Supplementary

Table 3). The mean value of three technical replicates was used for analysis. mRNA levels were normalized to *GAPDH* levels, and comparisons were made using the $\Delta\Delta$Ct method.

#### Quantification of MUC5AC-positive cells

MUC5AC-positive HBECs were quantified as described previously[2]. Either anti-MUC5AC-DL488 or anti-MUC5AC-DL405 (45M1, mouse $IgG_1$; Novus Biologicals, Centennial, CO) antibody was used. Flow cytometry (FACS Canto II, BD Biosciences, San Jose, CA) was performed at UCSF Laboratory for Cell Analysis, and data analysis was done with FlowJo (FlowJo LLC, Ashland, OR). The threshold for MUC5AC-positive cells was established based on staining of IL-13-treated cells with an isotype control antibody (<0.5% of isotype control-stained cells exceeded the threshold).

#### Immunofluorescence staining

Paraffin-embedded microscopic sections and whole mounts of HBEC culture Transwell inserts were prepared as described previously[27]. Immunofluorescence staining was done with following primary antibodies: mouse monoclonal anti-MUC5AC (45M1; Thermo Fisher Scientific; 1:200), rabbit polyclonal anti-MUC5B (H-300, sc-20119; Santa Cruz Biotechnology, Dallas, TX; 1:200), and mouse monoclonal anti acetylated alpha tubulin (6-11B-1, sc-23950; Santa Cruz Biotechnology; 1:200). After washing, slides were incubated with appropriate secondary antibodies (Rhodamine goat anti-mouse, Alex Fluor 647 goat anti-rabbit, and Alexa Fluor 488 goat anti-mouse; Jackson

ImmunoResearch Laboratories) at 1:200 dilution for 1 h. 4′,6-diamidinio-2-phenylindole (DAPI, 1:1000) was used to stain nuclei. Immunofluorescence images were acquired using a Yokagawa CSU22 spinning-disk confocal microscope connected to a Nikon Ti-E at Nikon Imaging Center and Center for Advanced Light Microscopy at UCSF. Slides were placed on the microscope stage and fluorescence and brightfield images were acquired using a 20X objective. Identical acquisition settings were used throughout each experiment. Images were processed using Fiji.

**Measurement of mucociliary transport**

Mucociliary transport was measured as described previously[2,27]. Briefly, 2-μm yellow-green (505/515) fluorescent microspheres (Thermo Fisher Scientific) were applied to the apical surface of HBEC cultures with intact mucus gels and allowed to disperse for 5 min. Cultures were then transferred to an optical cell dish (MatTek, Ashland, MA) and placed on the stage of a spinning disc confocal microscope under a dry 10X objective at 37 °C with perfluorocarbon (Sigma-Aldrich). Transport of fluorescent microspheres was imaged in the plane of the gel by recording sequential images every 1 s over 1 min. Images were analyzed using the TrackMate plugin in Fiji. Median microsphere speed was determined for each of three fields from one well per condition for each of three donors.

**Lentiviral GFP-based enhancer reporter assay**

Enhancer test sequences (Supplementary Table 4) were PCR-amplified from genomic DNA of BEAS-2B cells or synthesized (IDT, Newark, NJ) and cloned into XbaI and SbfI sites of the lentiviral GFP-based enhancer reporter plasmid pLS-mP[5] using the Quick Ligation Kit or NEBuilder HIFI Assembly (New England Biolabs, Ipswich, MA), respectively. HBECs were seeded at ~25% confluence in HPC-coated Transwell inserts (Corning, Corning, NY) and transduced with lentivirus. Transduced cells were maintained in ALI medium with 10 mM Y-27632 until 100% confluence and cultured at ALI for 23 d. Where indicated, PneumaCult-ALI Medium (STEMCELL Technologies, Vancouver, Canada) was used during the ALI culture, IL-13 (10 ng/mL) or IL-1$\beta$ (10 ng/mL) were added to the culture medium for the final 7 d of the culture period. Cells were harvested, fixed in 4% PFA for 10 min at 4°C, washed with PBS, and resuspended in eBioScience Flow Cytometry Staining Buffer (Thermo Fisher Scientific). GFP level was assessed by flow cytometry (FACSCanto II; BD Biosciences, San Jose, CA). In experiments assessing GFP levels in different subsets of HBECs, cells were blocked with 5% normal goat serum (Jackson ImmunoResearch Laboratories) and either stained with anti-NGFR/CD271-APC (ME20.4, $mIgG_1$; BioLegend, San Diego, CA) and anti-CEACAM6/CD66c-BV421 (B6.2, $mIgG_1$; BD Biosciences, San Jose, CA) antibodies to separate basal and secretory cells, or permeablized with 0.2% Saponin (Sigma-Aldrich, St. Louis, MO) and stained with anti-acetylated-$\alpha$-tubulin-AF647 (6-11B-1, $mIgG_{2b}$; Santa Cruz Biotechnology, Dallas, TX) antibody to separate ciliated cells[28]. In experiments with undifferentiated HBECs, BEAS-2B, and A549 cells, cells were seeded at ~25% confluence in HPC-coated or intact 12-well plates, transduced with lentivirus, and maintained in appropriate growth medium for 6 d post-transduction. Where indicated, IL-13 (10 ng/mL) was added to the culture medium for the final 4 d.

**Saturation mutagenesis MPRA**

Saturation mutagenesis MPRA was done as described previously[29] with minor modifications. All 600 possible single nucleotide variants of SPDEFe(1-200) were designed in addition to the wild-type, reference sequence and control sequences which included scrambled, SV40e, and IGHEe sequences, resulting in total of 638 sequences. A pool of 230-nt oligos containing each of these 200-nt sequences flanked by 15-nt primer recognition sequences was synthesized (SurePrint Oligonucleotide Libraries; Agilent Technologies, Santa Clara, CA), amplified, and cloned into the MPRA plasmid. Briefly, inserts were amplified and cloned upstream of a minimal promoter driving an *eGFP* reporter transcript with a barcode (BC). We aimed for ~50 BCs per sequence when constructing the plasmid library. Each insert was associated with a set of BCs using paired-end customized NGS (see Gordon et al. Step 83[30]). All MPRA-related sequencing was performed using an Illumina NextSeq 500 by the DNA Technologies & Expression Analysis Core Laboratory at UC Davis. 250,000 cells from each of two HBEC donors were transduced at a multiplicity of infection of 4, passaged to HPC-coated Transwell inserts, maintained in ALI medium with 10 mM Y-27632 until 100% confluence, and cultured at ALI for 23 d. Where indicated, IL-13 (10 ng/mL) was added to the culture medium for the final 7 d of culture.

~5,000,000 cells from each donor was harvested for each condition. Both genomic DNA and RNA were extracted using the AllPrep DNA/RNA mini kit (QIAGEN), and sequencing libraries were generated using a custom library preparation protocol for DNA and RNA extracts. DNA and RNA libraries were sequenced as specified in Gordon et al., Step 161[30]. MPRAflow (https://mpraflow.readthedocs.io/en/latest/quickstart.html) was used for association and count analysis. Barcode association was done using an amended association workflow, where we filtered for NM=0, mapq>0, and baseq>30. BCs were then filtered for coverage >3 and uniquely mapped to the enhancer sequence. A linear model

log2 RNA count ~ log2 DNA count + N+ intercept

was fitted for each variant enhancer sequence against the wildtype enhancer sequence, where N=1 for wildtype and N=0 for the variant sequence. Multiple testing adjustment was done where *p*-values were adjusted by FDR. Enhancer sequences with FDR < 0.05 were considered significant.

**CRISPR-based gene targeting**

Gene targeting via CRISPR in HBECs was done as described previously[2]. gRNA sequences are listed in Supplementary Table 2. Synthetic sgRNAs from Synthego (Redwood City, CA) and rCas9 from MacroLabs (Berkeley, CA) were used; nucleofections were performed with the 4D-Nucleofector System (program DC-100; Lonza). Transfected cells were subsequently transduced with lentivirus containing GFP-based enhancer reporter constructs. Cells were then passaged to Transwell inserts for culture at ALI for 23 d. Where indicated, IL-13 (10 ng/mL) was added for the final 7 d of culture.

**KLF5 CUT&Tag**

HBECs from three donors (10-75, 13-32, and 14-30) cultured at ALI for 23 d. Where indicated, IL-13 (10 ng/mL) was added for the final 7 d of culture. KLF5 CUT&Tag was conducted using a Hyperactive pA-Tn5 In-Situ ChIP Library Prep Kit for Illumina (Vazyme Biotech Co., Nanjing, China) with minor modifications to the manufacturer's protocol as follows. Anti-KLF5 (ab137676, rabbit IgG; Abcam) polyclonal antibodies were used, and normal rabbit IgG was used as a negative control (NI01; Sigma-Aldrich). No secondary antibody was used. The starting cell amount was 500,000 cells, and primary antibody incubation was done overnight. Post-amplification, magnetic bead-based clean-up (HighPrep PCR Clean-up System; MAGBIO Genomics, Gaithersburg, MD) was conducted twice for each clean-up step. We used a small amount of the tagmented and pre-amplified library to set the final number of cycles needed for final library amplification. The cycle threshold (Ct) for which the library had reached 1/3 of its maximal fluorescence (Rn) was set independently for each sample. Libraries were sequenced (paired end 150 bp) using NextSeq (DNA Technologies & Expression Analysis Core Laboratory at UC Davis) and NovaSeq SP 300 (Center for Advanced Technology at UCSF) sequencers. Reads were aligned with STAR with reference hg38. Peaks were called with MACS2 (parameters -f BAM --keep-dup 1  -p 0.01 –nomodel). Overlaps between reproducible peaks were found using IDR. The union of the Cut&Tag

peaks with IDR score > 540 (or reproducibility < 0.05) across the same donor and treatment conditions were combined into one file and used in subsequent analyses. HOMER Motif analysis was performed with parameter size –given. Intersections of peaks from two technical replicates per donor and three donors were used in the differentially modulated peaks analysis. To find peaks modulated by IL-13 stimulation, R package DESEQ2 was used (R version 4.0.1, DESeq2 version 1.28.1). Samtools was used to extract counts from all three donors across two conditions, obtained from the merged bam files of for each donor/treatment condition. Normalization through median of ratios was performed, and a negative binomial GLM was fitted to the count data. Peaks with $p$-value < 0.1 were called as significantly upregulated or downregulated peaks.

**Enhancer-driven CRISPRi targeting**

Enhancer sequences (Supplementary Table 4) were PCR-amplified from genomic DNA from BEAS-2B cells or synthesized (IDT) and cloned into XbaI and SbfI sites of the mP-KRAB-dCas9_EF1a-BSD plasmid using the Quick Ligation Kit or NEBuilder HIFI Assembly. HBECs were initially transduced with enhancer-mP-KRAB-dCas9 lentivirus and maintained in BEGM with 10 μM Y-27632 and 10 μg/mL blasticidin (Thermo Fisher Scientific) for 3 d for selection. Cells were subsequently transduced with sgRNA lentivirus and maintained in BEGM with 10 μM Y-27632, 10 μg/mL blasticidin, and 1 μg/mL puromycin (Thermo Fisher Scientific) for 3 d for selection. Transduced and selected cells were then passaged to Transwell inserts for culture at ALI for 23 d. Where indicated, IL-13 (10 ng/mL) was added for the final 7 d of culture.

**Supplementary Data**

**Supplementary Data 1.** Bulk RNA-seq of HBECs without and with IFN-$\alpha$, IFN-$\gamma$, IL-17, IL-13, or IFN-$\alpha$ and IL-13 stimulation.

**Supplementary Data 2.** Single cell RNA-seq of three major HBEC subsets (basal, ciliated, and secretory) without and with IFN-$\alpha$, IL-17, or IL-13 stimulation.

**Supplementary Data 3.** Gene set enrichment analysis of IFN-$\alpha$, IL-17, or IL-13 response in three major HBEC subsets (basal, ciliated, and secretory).

**Supplementary Data 4.** Saturation mutagenesis MPRA of SPDEFe(1-200).

#### Supplementary Tables

**Supplementary Table 1.** Statistical comparisons for Fig. 3f by one-way ANOVA Tukey's post-test.

| Condition[a] | 1 | 2 | 3 | 4 | 5 | 6 | 7 | 8 | 9 | 10 | 11 | 12 | 13 | 14 | 15 | 16 | 17 | 18 |
|---|---|---|---|---|---|---|---|---|---|---|---|---|---|---|---|---|---|---|
| 1 | | ns | **** | **** | ** | *** | **** | **** | **** | **** | **** | *** | **** | **** | **** | **** | **** | **** |
| 2 | | | **** | **** | ** | ** | **** | **** | **** | **** | **** | *** | **** | **** | **** | **** | **** | **** |
| 3 | | | | ns | **** | **** | ns | ns | ns | ** | **** | **** | **** | **** | ns | ns | ns | ns |
| 4 | | | | | **** | **** | ns | ns | ns | ** | **** | **** | **** | **** | ns | ns | ns | ns |
| 5 | | | | | | ns | **** | **** | **** | **** | ns | ns | ns | ns | **** | **** | **** | **** |
| 6 | | | | | | | **** | **** | **** | **** | ns | ns | ns | ns | **** | **** | **** | **** |
| 7 | | | | | | | | ns | ns | ** | **** | **** | **** | **** | ns | ns | ns | ns |
| 8 | | | | | | | | | ns | ns | **** | **** | **** | **** | ns | ns | ns | ns |
| 9 | | | | | | | | | | ns | **** | **** | **** | **** | ns | ns | ns | ns |
| 10 | | | | | | | | | | | **** | **** | **** | **** | * | ns | ns | **** |
| 11 | | | | | | | | | | | | ns | ns | ns | **** | **** | **** | **** |
| 12 | | | | | | | | | | | | | ns | ns | **** | **** | **** | **** |
| 13 | | | | | | | | | | | | | | ns | **** | **** | **** | **** |
| 14 | | | | | | | | | | | | | | | **** | **** | **** | **** |
| 15 | | | | | | | | | | | | | | | | ns | ns | ns |
| 16 | | | | | | | | | | | | | | | | | ns | ns |
| 17 | | | | | | | | | | | | | | | | | | ns |
| 18 | | | | | | | | | | | | | | | | | | |

[a]1, NT-1(-); 2, NT-2(-); 3, NT-1; 4, NT-2; 5, SPDEF-TSS(+34); 6, SPDEF-TSS(+45); 7, SPDEFe-vicinity(+4091); 8, SPDEFe-vicinity(+4071); 9, SPDEFe-vicinity(+2089); 10, SPDEFe-vicinity(+2032); 11, SPDEFe(+87); 12, SPDEFe(+53); 13, SPDEFe(-11); 14, SPDEFe(-164); 15, SPDEFe-vicinity(+-2124); 16, SPDEFe-vicinity(-2157); 17, SPDEFe-vicinity(-4320); and 18, SPDEFe-vicinity(-4059). ns, not significant; *, $p < 0.05$; **, $p < 0.01$; ***, $p < 0.001$; and ****, $p < 0.0001$.

**Supplementary Table 2.** gRNA sequences.[b]

| Name[a] | Sequence (5′-3′) |
|---|---|
| NT-1 | GACGACTAGTTAGGCGTGTA |
| NT-2 | GGCCAAACGTGCCCTGACGG |
| SPDEF-TSS(+1) | ACAGCCGCGAGATGAAGAGT |
| SPDEF-TSS(+34) | GCAGAGTGCAGGAATGTGCT |
| SPDEF-TSS(+45) | GTGTGGACACGGCAGAGTGC |
| SPDEF-TSS(+232) | GCACTCAGGTTGGCCACTGG |
| MUC5AC-TSS(+36) | AACACTCATTGTGTGGACGG |
| MUC5AC-TSS(+117) | CTGGCCTGCACCCGGCATAC |
| MUC5AC-TSS(+134) | AGGACCAGTAGAGCGGCCAG |
| MUC5AC-TSS(+172) | TGGGTGGTGCGGTACTGAGT |
| SPDEFe(-164) | TGGCCTGAGCCATTCCCACG |
| SPDEFe(-11) | GGCCTTCTCAGGAACAGGGT |
| SPDEFe(+53) | ATGGTTGGGTCACAGCAAGC |
| SPDEFe(+87) | ACCATTTCCCCGACATCACC |
| SPDEFe-vicinity(-4320) | TTGCCATACAAAACCTCGGC |
| SPDEFe-vicinity(-4059) | CGATGTGGTCTGCAATCCAG |
| SPDEFe-vicinity(-2157) | CTGCCAGGTACCCACGCCAC |
| SPDEFe-vicinity(-2124) | GCAGCAGTCTGGGCCTAATG |
| SPDEFe-vicinity(+2032) | GCTGGCAGCGTGGATATACA |
| SPDEFe-vicinity(+2089) | GATAACCTCAACCAGTGCTG |
| SPDEFe-vicinity(+4071) | GGTACTTACACAGGTCCTTG |
| SPDEFe-vicinity(+4091) | AAGGACCTGTGTAAGTACCA |
| STAT6 | AGGGAATGGCGCACCGTTTG |
| KLF5 | TGAAACGACTGCCCCTCCTC |

[b]*SPDEF* and *MUC5AC* promoter/TSS targeting gRNAs are numbered relative to their TSSs (chr6:34,556,333 and chr11:1,157,953). SPDEFe targeting and its neighboring gRNAs are numbered relative to the mid-point of the essential STAT6 motif within SPDEFe (chr6:34,586,431).

**Supplementary Table 3.** RT-qPCR primer sequences.

| Name | Sequence (5′-3′) |
|---|---|
| GAPDH.f | CACATCGCTCAGACACCAT |
| GAPDH.r | GGCAACAATATCCACTTTACCAG |
| SPDEF.f | AAGTTGGCACTGCAGCAGAC |
| SPDEF.r | GGGGATACGCTGCTCAGAC |
| FOXA3.f | TGGCCGAGTGGAGCTACTAC |
| FOXA3.r | GGATTCAGGGTCATGTAGGAGT |
| MUC5AC.f | CAACATCAGGAACAGCTTCGA |
| MUC5AC.r | GAGCACCAGTGCTGAGCAT |
| SCGB1A1.f | GCTGAAGAAGCTGGTGGACAC |
| SCGB1A1.r | TGCTAATTACACAGTGAGCTTTGG |
| POSTN.f | GACCGTGTGCTTACACAAATTG |
| POSTN.r | AAGTGACCGTCTCTTCCAAGG |
| KRT5.f | AGCAGTGGTACGCTTGTTGATT |
| KRT5.r | GCCTGGACTCAGAGCTGAGAA |
| TUBB4B.f | ATCTTCCGGCCGGACAAC |
| TUBB4B.r | CATCCAGCACCGAGTCCA |
| HMGA1.f | ACAAGGGTGCTGCCAAGAC |
| HMGA1.r | CCGAGGACTCCTGCGAGAT |
| NUDT3.f | TGTGTTTCCGCAGCGAGA |
| NUDT3.r | CTCCTCCAGGGACAATCCAT |
| ILRUN.f | CCTTCTTCCTGGACATGACCA |
| ILRUN.r | GGACATAGAGGGCACACTGATG |
| SNRPC.f | GAGCAGGCTCAGAGCCTGAT |
| SNRPC.r | GGGAGGTGGTATCATCGCC |
| FKBP5.f | AAACATCCTTCCACCACAGC |
| FKBP5.r | GAAGATGGAGGCATTATCCG |
| dCas9.f | CAAAGTGATGAAGCAGCTGAAG |
| dCas9.r | GTCGGACTTCAGGAAATCCA |

**Supplementary Table 4.** Enhancer test sequences used in lentiviral GFP-based reporter assays and enhancer-driven CRISPRi.

| Name | Sequence (5′-3′) |
|---|---|
| SV40e | GCGCAGCACCATGGCCTGAAATAACCTCTGAAAGAGGA<br>ACTTGGTTAGGTACCTTCTGAGGCGGAAAGAACCAGCT<br>GTGGAATGTGTGTCAGTTAGGGTGTGGAAAGTCCCCAG<br>GCTCCCCAGCAGGCAGAAGTATGCAAAGCATGCATCTC<br>AATTAGTCAGCAACCAGGTGTGGAAAGTCCCCAGGCTC<br>CCCAGCAGGCAGAAGTATGCAAAGCATGCATCTCAATT<br>AGTCAGCAACCATAGTCCCGCCCCTAACTCCGCCCATC<br>CCGCCCCTAACTCCGCCCAGTTCCGCCCATTCTCCGCC<br>CCATGGCTG |
| IGHEe | CGCTGTTGCTCAATCGACTTCCCAAGAACAGCGCTGTT<br>GCTCAATCGACTTCCCAAGAACAG |
| SPDEFe | GGTGATGTCGGGGAAATGGTTGGGTCACAGCAAGCAGG<br>TGGCCTTCTCAGGAACCTGCGTGAGAAGGGCCTGAGGG<br>AGAGGCCTTCTCAGGAACAGGGTGGGAAGCACCCAGAC<br>ACAACCTGAGCTGAGGCACAGAGGAGACTTGAGAGTGC<br>CTGTTTGCTCTGCCAGAAGGGTTGGGCCCAGACCACTG<br>CAATTGCTTCCCTGCCATCTCCCAAACTCAGCTGAGAG<br>CCCCCTCCCTGTCACCCAGCCCCCGTGGGAATGGCTCA<br>GGCCAGGAGCTACCATCAAACTAGCTCCTTTCTCCCAG<br>GGCCTGGTGACTCCTTGACAGGTTACAGCCCCAAAGAC<br>AAACAGATTCTGCCAAGCAAATAACTGTTCAGCAGTGG<br>GGGTGTTGGGGAAGGCCTAGGAGTGGGTCTGAAGCTGC<br>CAGAAGAAGACCCAGGTGAAGTGGGGGGTGGGCTCCCC<br>TGCAGAGAACAAACAGCTGTGGGCACCTGGAAAGAAGC<br>ACGATGCAGGTACACTGTCACGGGTGAGGGCACTCACC<br>CCAGGGGTCTGTGGGCCAGTCTCCCAGGCTCTTTCACT<br>CTGGGCTCTGTATCAGGCT |
| SPDEFe(1-147) | GGTGATGTCGGGGAAATGGTTGGGTCACAGCAAGCAGG<br>TGGCCTTCTCAGGAACCTGCGTGAGAAGGGCCTGAGGG<br>AGAGGCCTTCTCAGGAACAGGGTGGGAAGCACCCAGAC<br>ACAACCTGAGCTGAGGCACAGAGGAGACTTGAG |
| SPDEFe(70-226) | CTGAGGGAGAGGCCTTCTCAGGAACAGGGTGGGAAGCA<br>CCCAGACACAACCTGAGCTGAGGCACAGAGGAGACTTG<br>AGAGTGCCTGTTTGCTCTGCCAGAAGGGTTGGGCCCAG<br>ACCACTGCAATTGCTTCCCTGCCATCTCCCAAACTCAG<br>CTGAG |
| SPDEFe(140-294) | GACTTGAGAGTGCCTGTTTGCTCTGCCAGAAGGGTTGG<br>GCCCAGACCACTGCAATTGCTTCCCTGCCATCTCCCAA<br>ACTCAGCTGAGAGCCCCCTCCCTGTCACCCAGCCCCCG<br>TGGGAATGGCTCAGGCCAGGAGCTACCATCAAACTAGC<br>TCC |
| SPDEFe(213-361) | CAAACTCAGCTGAGAGCCCCCTCCCTGTCACCCAGCCC<br>CCGTGGGAATGGCTCAGGCCAGGAGCTACCATCAAACT<br>AGCTCCTTTCTCCCAGGGCCTGGTGACTCCTTGACAGG<br>TTACAGCCCCAAAGACAAACAGATTCTGCCAAGCA |
| SPDEFe(288-442) | TAGCTCCTTTCTCCCAGGGCCTGGTGACTCCTTGACAG<br>GTTACAGCCCCAAAGACAAACAGATTCTGCCAAGCAAA<br>TAACTGTTCAGCAGTGGGGGTGTTGGGGAAGGCCTAGG |

| | |
|---|---|
| | AGTGGGTCTGAAGCTGCCAGAAGAAGACCCAGGTGAAG TGG |
| SPDEFe(363-515) | ATAACTGTTCAGCAGTGGGGGTGTTGGGGAAGGCCTAG GAGTGGGTCTGAAGCTGCCAGAAGAAGACCCAGGTGAA GTGGGGGGTGGGCTCCCCTGCAGAGAACAAACAGCTGT GGGCACCTGGAAAGAAGCACGATGCAGGTACACTGTCA C |
| SPDEFe(450-589) | GCTCCCCTGCAGAGAACAAACAGCTGTGGGCACCTGGA AAGAAGCACGATGCAGGTACACTGTCACGGGTGAGGGC ACTCACCCCAGGGGTCTGTGGGCCAGTCTCCCAGGCTC TTTCACTCTGGGCTCTGTATCAGGCT |
| SPDEFe(1-147)_1stat6 | GGTGATGTCGGGGAAATGGTTGGGTCACAGCAAGCAGG TGGCCCAGGTCCTGCCCTGCGTGAGAAGGGCCTGAGGG AGAGGCCTTCTCAGGAACAGGGTGGGAAGCACCCAGAC ACAACCTGAGCTGAGGCACAGAGGAGACTTGAG |
| SPDEFe(1-147)_2stat6 | GGTGATGTCGGGGAAATGGTTGGGTCACAGCAAGCAGG TGGCCTTCTCAGGAACCTGCGTGAGAAGGGCCTGAGGG AGAGGCCCAGGTCCTGCCAGGGTGGGAAGCACCCAGAC ACAACCTGAGCTGAGGCACAGAGGAGACTTGAG |
| SPDEFe(1-147)_1&2stat6 | GGTGATGTCGGGGAAATGGTTGGGTCACAGCAAGCAGG TGGCCCAGGTCCTGCCCTGCGTGAGAAGGGCCTGAGGG AGAGGCCCAGGTCCTGCCAGGGTGGGAAGCACCCAGAC ACAACCTGAGCTGAGGCACAGAGGAGACTTGAG |
| SPDEFe(1-147)_1klf5 | GGTGATGTCGGGGAAATGGTTGGGTCACAGCAAGCAGG TGGCCTTCTCAGGAACCTGCGTGAGAAGGGCCTGATGC GATTACCTTCTCAGGAACAGGGTGGGAAGCACCCAGAC ACAACCTGAGCTGAGGCACAGAGGAGACTTGAG |
| SPDEFe(1-147)_2klf5 | GGTGATGTCGGGGAAATGGTTGGGTCACAGCAAGCAGG TGGCCTTCTCAGGAACCTGCGTGAGAAGGGCCTGAGGG AGAGGCCTTCTCAGGAACATGCGATTAAGCACCCAGAC ACAACCTGAGCTGAGGCACAGAGGAGACTTGAG |
| SPDEFe(1-147)_1&2klf5 | GGTGATGTCGGGGAAATGGTTGGGTCACAGCAAGCAGG TGGCCTTCTCAGGAACCTGCGTGAGAAGGGCCTGATGC GATTACCTTCTCAGGAACATGCGATTAAGCACCCAGAC ACAACCTGAGCTGAGGCACAGAGGAGACTTGAG |
| SPDEFe(1-147)x2 | GGTGATGTCGGGGAAATGGTTGGGTCACAGCAAGCAGG TGGCCTTCTCAGGAACCTGCGTGAGAAGGGCCTGAGGG AGAGGCCTTCTCAGGAACAGGGTGGGAAGCACCCAGAC ACAACCTGAGCTGAGGCACAGAGGAGACTTGAGGGTGA TGTCGGGGAAATGGTTGGGTCACAGCAAGCAGGTGGCC TTCTCAGGAACCTGCGTGAGAAGGGCCTGAGGGAGAGG CCTTCTCAGGAACAGGGTGGGAAGCACCCAGACACAAC CTGAGCTGAGGCACAGAGGAGACTTGAG |
| SPDEFe(1-147)x3 | GGTGATGTCGGGGAAATGGTTGGGTCACAGCAAGCAGG TGGCCTTCTCAGGAACCTGCGTGAGAAGGGCCTGAGGG AGAGGCCTTCTCAGGAACAGGGTGGGAAGCACCCAGAC ACAACCTGAGCTGAGGCACAGAGGAGACTTGAGGGTGA TGTCGGGGAAATGGTTGGGTCACAGCAAGCAGGTGGCC TTCTCAGGAACCTGCGTGAGAAGGGCCTGAGGGAGAGG CCTTCTCAGGAACAGGGTGGGAAGCACCCAGACACAAC CTGAGCTGAGGCACAGAGGAGACTTGAGGGTGATGTCG GGGAAATGGTTGGGTCACAGCAAGCAGGTGGCCTTCTC AGGAACCTGCGTGAGAAGGGCCTGAGGGAGAGGCCTTC |

| | |
|---|---|
| | TCAGGAACAGGGTGGGAAGCACCCAGACACAACCTGAG<br>CTGAGGCACAGAGGAGACTTGAG |

### Supplementary Figure 1

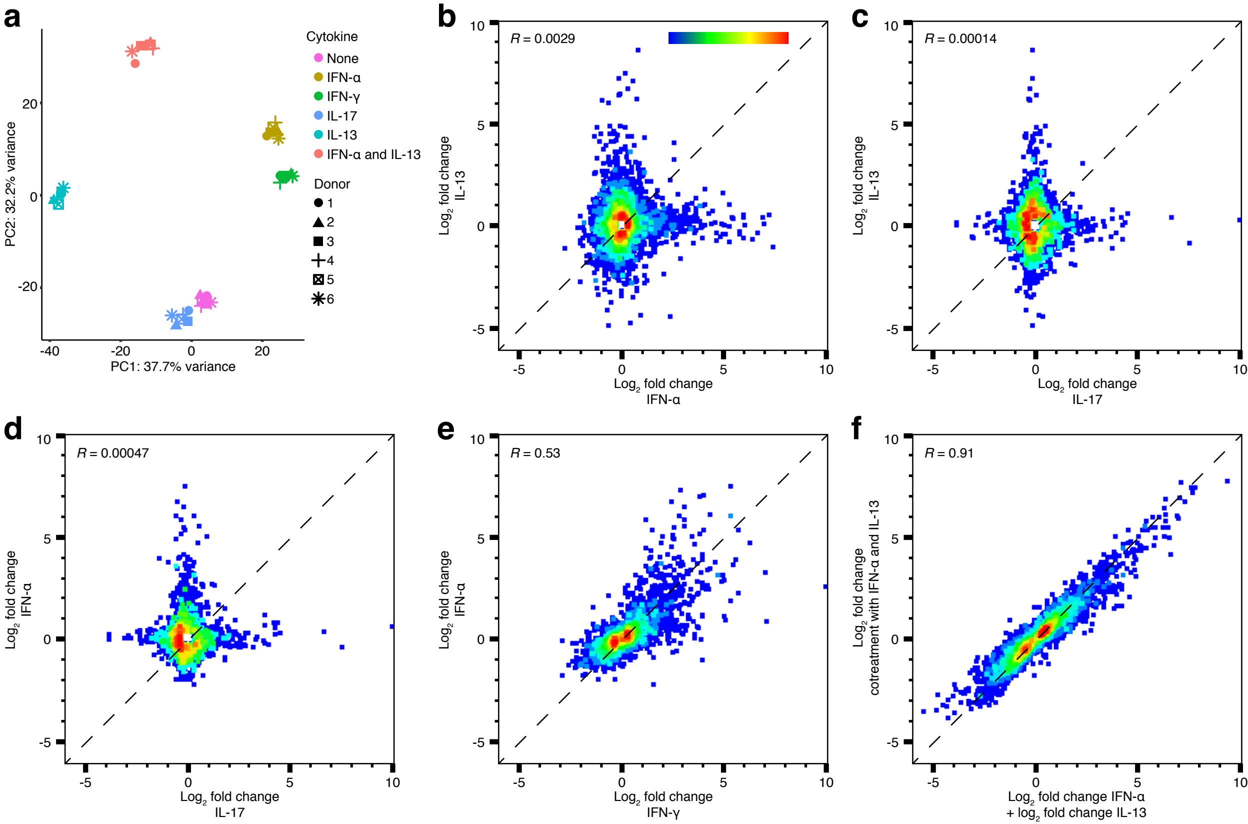

### Supplementary Figure 2

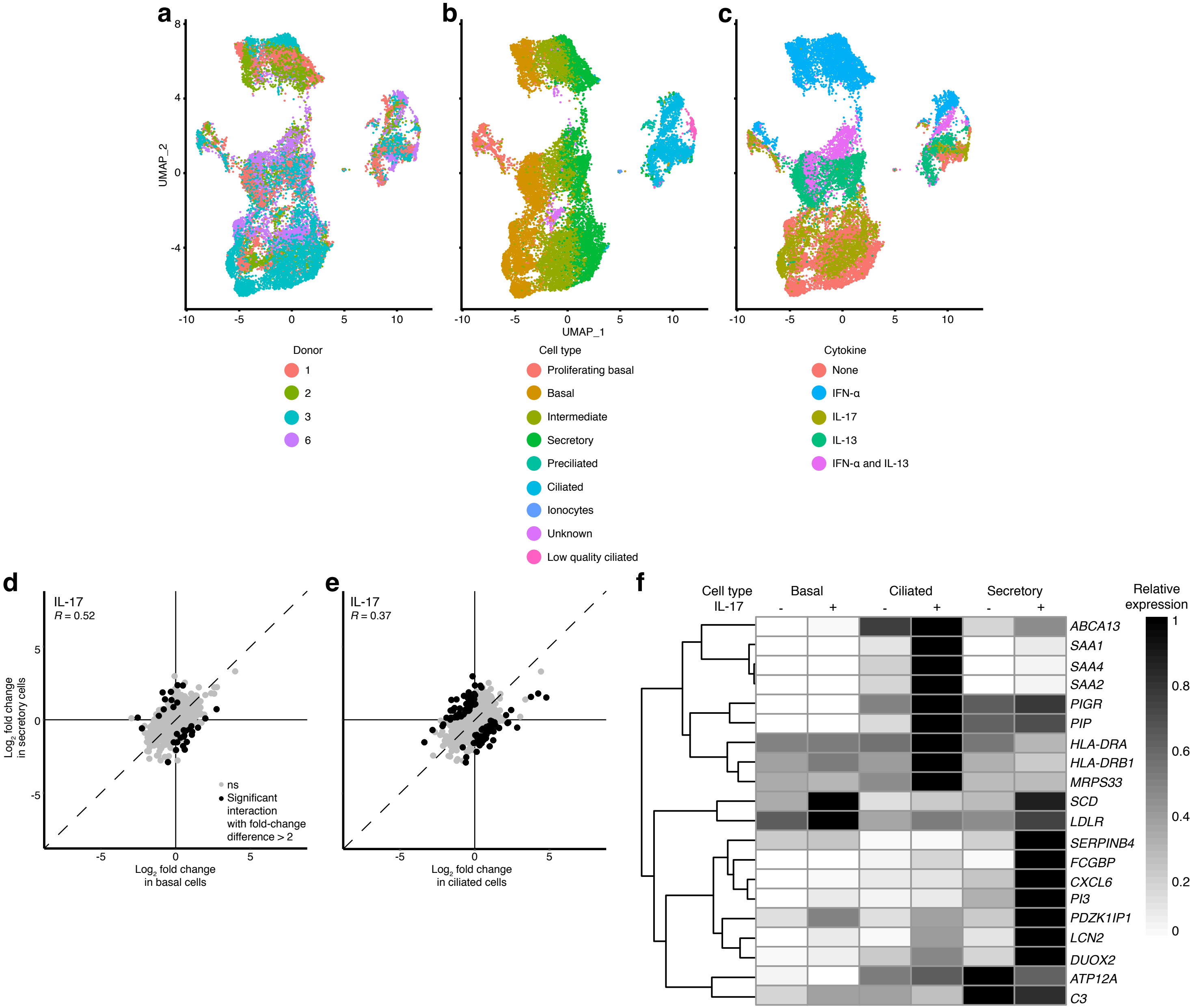

### Supplementary Figure 3

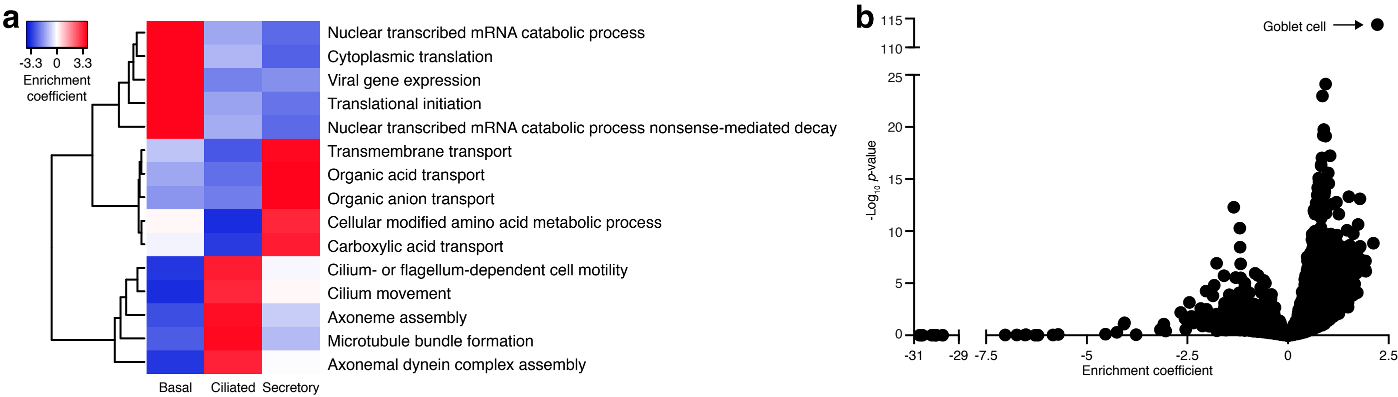

### Supplementary Figure 4

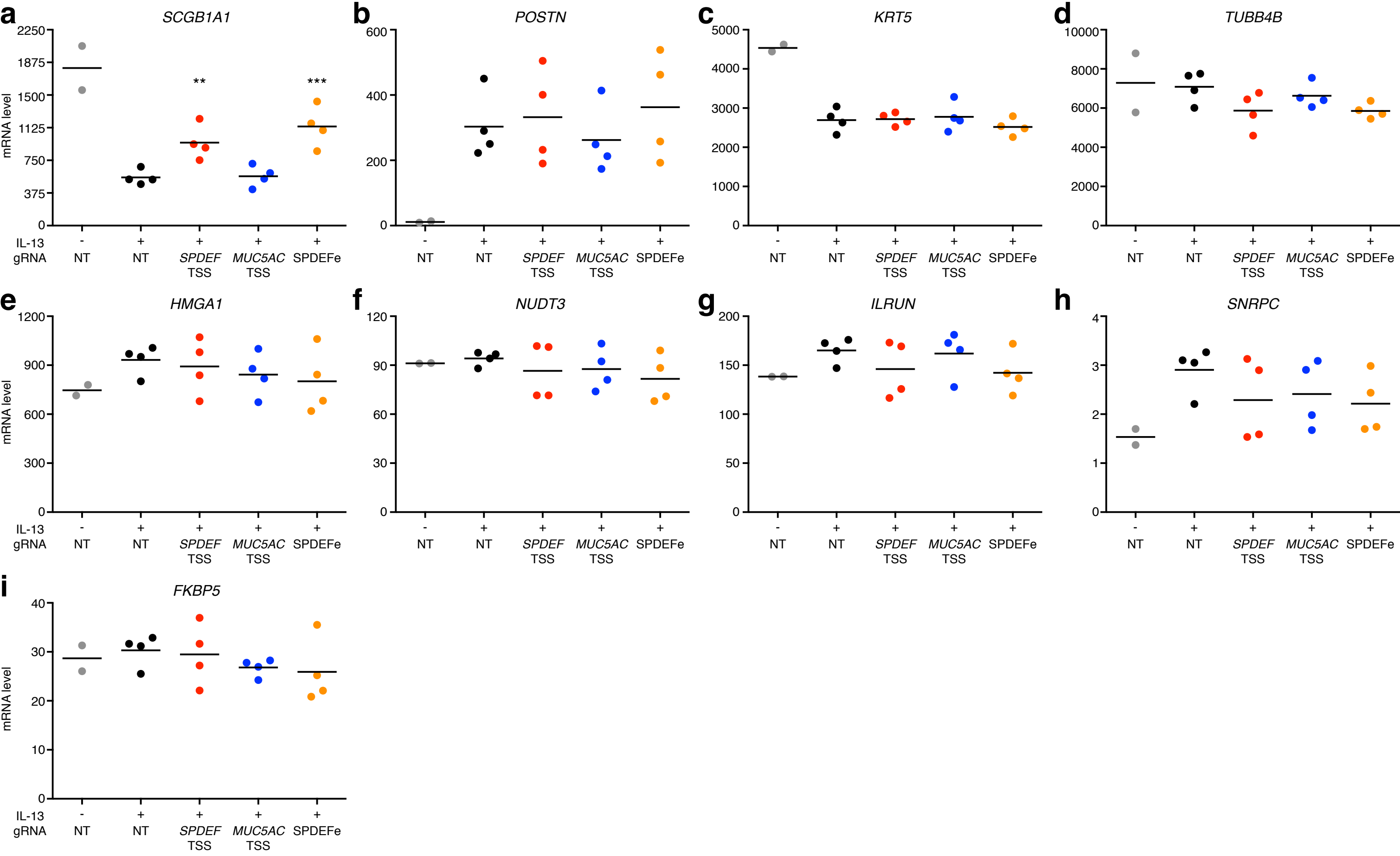

### Supplementary Figure 5

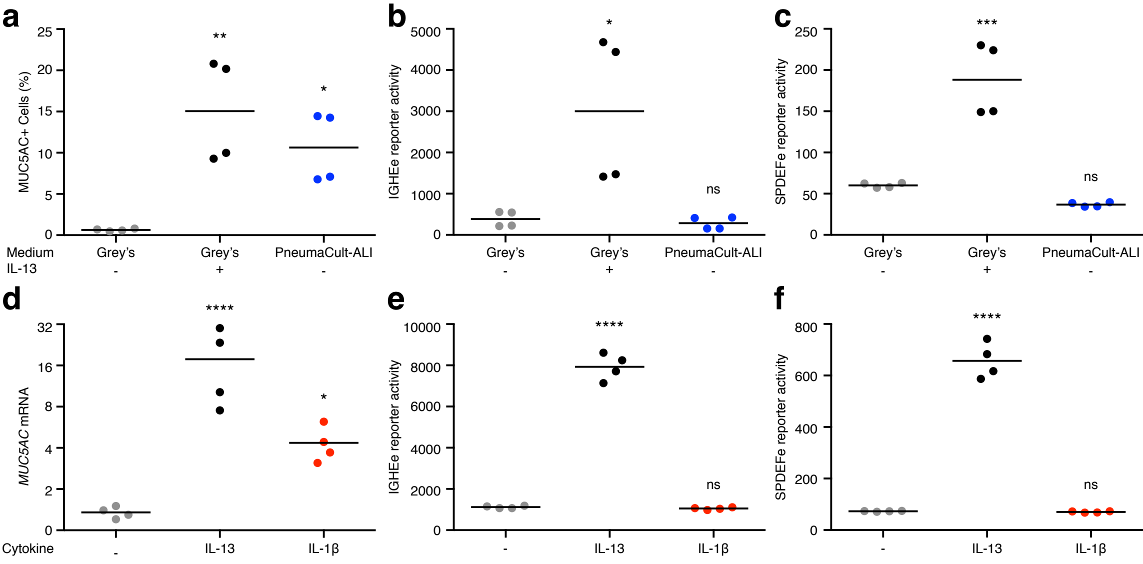

### Supplementary Figure 6

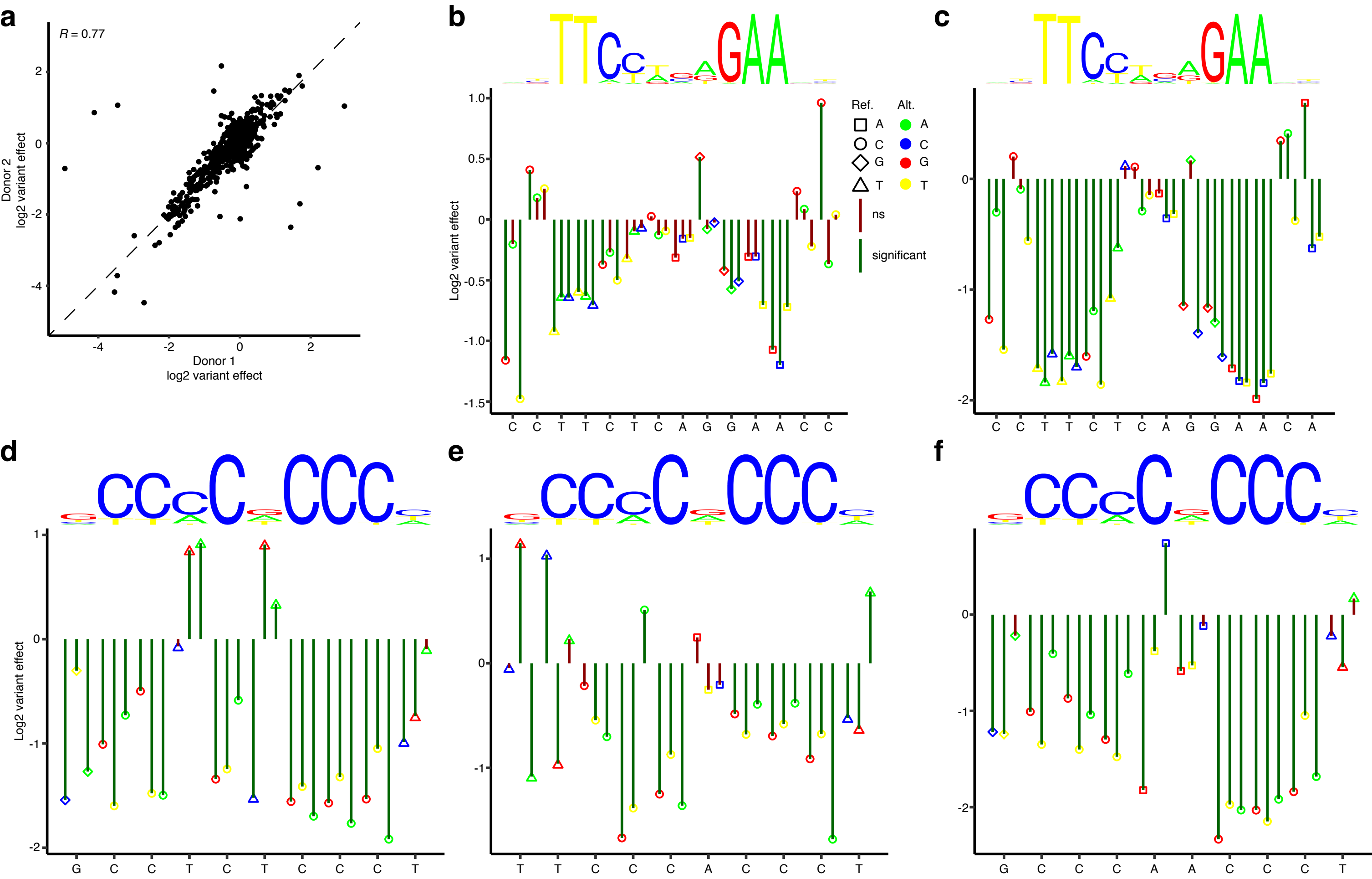

#### Supplementary Figure Legends

**Supplementary Figure 1.** Transcriptional effects of asthma-associated cytokines on HBECs as determined by bulk RNA-seq. **a**, Principal Component Analysis (PCA) of bulk RNA-seq results from cultures from six HBEC donors that were unstimulated or stimulated with IFN-$\alpha$, IFN-$\gamma$, IL-17, IL-13, or the combination of IFN-$\alpha$ and IL-13. **b**-**e**, Pairwise comparisons of fold changes in gene expression induced by IFN-$\alpha$ compared with IL-13 (**b**), IL-17 compared with IL-13 (**c**), IL-17 compared with IFN-$\alpha$ (**d**), and IFN-$\gamma$ compared with IFN-$\alpha$ (**e**). **f**, Pairwise comparison of sum of fold changes induced by individual stimulation with IFN-$\alpha$ and IL-13 compared with fold change induced by combined stimulation with IFN-$\alpha$ and IL-13. *R*, Pearson correlation coefficient ($p < 0.0001$).

**Supplementary Figure 2.** Transcriptional effects of asthma-associated cytokines on HBECs as determined by scRNA-seq. **a**-**c**, UMAP analysis of scRNA-seq results from cultures from four HBEC donors that were unstimulated or stimulated with IFN-$\alpha$, IFN-$\gamma$, IL-17, IL-13, or the combination of IFN-$\alpha$ and IL-13. Each point represents a single cell, and points are colored by donor (**a**), airway epithelial cell type (**b**), and cytokine treatment (**c**). **d** and **e**, Effects of IL-17 on secretory cells compared with basal cells (**d**) or ciliated cells (**e**). Each point represents a single gene. Genes that were regulated differently between cell types are in black ($\log_2$ fold-change difference > 1 and FDR < 0.1 for interactions between cell type and cytokine effect), and all other genes are in grey. *R*, Pearson correlation coefficient ($p < 2.2 \times 10^{-16}$ for both comparisons). **f**,

Relative expression of the 20 genes that were most strongly regulated by IL-17. These 20 genes were selected based on the highest absolute fold change in any of the three cell types.

**Supplementary Figure 3.** Gene set enrichment analysis of IL-13-induced cell type-selective responses in HBECs. To explore cell type-selective effects of IL-13, we performed gene set enrichment analysis of IL-13-induced responses in basal, ciliated, and secretory cells using our scRNA-seq data. **a**, Relative enrichment coefficients of the 5 most highly enriched GOBP gene sets in basal, ciliated, and secretory cells with IL-13 stimulation ($p < 10^{-5}$ in one cell type and enrichment coefficient < 0.5 in other cell types). **b**, The set of genes that were induced in secretory cells following IL-13 stimulation is highly enriched for a previously defined set of goblet cell genes[31]. Other points represent values for the full set of 2,555 GOBP gene sets.

**Supplementary Figure 4.** Effects of CRISPRi targeting SPDEFe on gene expression. These data were acquired as a part of the same experiment performed in Fig. 3b-e. HBECs were transduced with lentiviruses driving expression of *dCas9-KRAB* and either non-targeting (NT) control sgRNAs (grey/black) or sgRNAs targeting the *SPDEF* promoter (red), the *MUC5AC* promoter (blue), or SPDEFe (orange). After differentiation, cells were left unstimulated or stimulated with IL-13 for 7 days, as indicated. Gene expression was measured by qRT-PCR. We measured the expression of a gene encoding a club cell-secreted secretoglobin (*SCGB1A1*, **a**) which is suppressed by Th2 cytokines and is less in individuals with asthma than those without asthma[32]. As its name and alias (*CCSP*) indicate, a high level of *SCGB1A1* is a marker of precursor of goblet cells. An increase, proportionate to the decrease observed in *SPDEF* and *MUC5AC*, in expression *SCGB1A1* was observed when targeting SPDEFe or the *SPDEF* TSS. To exclude the possibility that targeting SPDEFe has non-specific effects on IL-13 responsiveness, we measured expression of periostin (*POSTN*, **b**), a gene that is frequently used as part of an IL-13-response signature in asthma. Targeting the *SPDEF* TSS, *MUC5AC* TSS, or SPDEFe had no effects on this IL-13-inducible gene as well as on the expression of the basal cell marker *KRT5* (**c**) and the ciliated cell marker *TUBB4B* (**d**). We also measured effects of SPDEFe gRNAs on the neighboring genes *HMGA1* (**e**), *NUDT3* (**f**), *ILRUN* (**g**), *SNRPC* (**h**), and *FKBP5* (**i**) and found no effects. Values are mRNA copy numbers per 1,000 copies of *GAPDH*. Each point corresponds to a different gRNA targeting the indicated region, tested separately in a single culture well from the same donor. **, $p < 0.01$ and ***, $p < 0.001$ for comparison with IL-13-stimulated HBECs with NT control sgRNAs by one-way ANOVA with Dunnett's post-test; all other differences were not statistically significant.

**Supplementary Figure 5.** Assessment SPDEFe activity in PneumaCult-ALI medium and with IL-1β stimulation. **a**-**c**, HBECs were transduced with IGHEe or SPDEFe lentiviral GFP reporter constructs and cultured at ALI. Cells were cultured in standard Grey's medium without cytokine stimulation (grey) or with IL-13 (black) for the last 7 d of culture, or in PneumaCult-ALI medium without cytokine stimulation (blue). Intracellular MUC5AC (**a**) and reporter activity (**b** and **c**) were determined by flow cytometry. Data were obtained from four donors. ns, not significant; *, $p < 0.05$; **, $p < 0.01$; and ***, *p* <

0.001 for comparison against Grey's medium with no stimulation by one-way ANOVA with Tukey's post-test. **d**-**f**, HBECs were transduced with IGHEe and SPDEFe lentiviral reporters and cultured in standard Grey's medium without cytokine stimulation (grey) or with IL-13 (black) or IL-1β (red) for the last 7 d of culture. Expression of *MUC5AC* (**d**) was measured by qRT-PCR. Values are mRNA copy numbers per 1,000 copies of *GAPDH*. Reporter activity (**e** and **f**) were determined by flow cytometry. Data were obtained from four donors. ns, not significant; *, $p < 0.05$; and ****, $p < 0.0001$ for comparison against unstimulated by one-way ANOVA Tukey's post-test.

**Supplementary Figure 6.** Saturation mutagenesis MPRA of bases 1-200 of SPDEFe. **a**, Comparison of effects of each single nucleotide substitution between two donors. *R*, Pearson correlation coefficient ($p < 2.2 \times 10^{-16}$). **b**-**f**, Effects of each single nucleotide substitutions (combined data from two donors) in two STAT6 motifs, 42-55 (**b**) and 82-95 (**c**), and three KLF5 motifs (shown in reverse complement), 82-73 (**d**), 104-95 (**e**), and 179-170 (**f**).
